# Adaptive stimulation of macropinocytosis overcomes aspartate limitation in cancer cells under hypoxia

**DOI:** 10.1101/2021.02.02.429407

**Authors:** Javier Garcia-Bermudez, Sheela Prasad, Lou Baudrier, Michael A. Badgley, Yuyang Liu, Konnor La, Mariluz Soula, Robert T. Williams, Norihiro Yamaguchi, Rosa F. Hwang, Laura J. Taylor, Elisa De Stanchina, Bety Rostandy, Hanan Alwaseem, Henrik Molina, Dafna Bar-Sagi, Kıvanç Birsoy

**Affiliations:** Laboratory of Metabolic Regulation and Genetics, The Rockefeller University, New York, NY, USA; Department of Biochemistry and Molecular Pharmacology, New York University School of Medicine, New York, NY, USA; Laboratory of Systems Cancer Biology, The Rockefeller University, New York, NY, USA; Department of Breast Surgical Oncology, Division of Surgery, MD Anderson Cancer Center, University of Texas, Houston, Texas; Antitumor Assessment Core Facility, Memorial Sloan Kettering Cancer Center, New York, NY, USA; The Proteomics Resource Center, The Rockefeller University, New York, NY, USA

## Abstract

Stress-adaptive mechanisms enable tumor cells to overcome metabolic constraints in nutrient and oxygen poor tumors. Aspartate is an endogenous metabolic limitation under hypoxic conditions, but the nature of the adaptive mechanisms that contribute to aspartate availability and hypoxic tumor growth are poorly understood. Here, using a combination of metabolomics and CRISPR-based genetic screens, we identify GOT2-catalyzed mitochondrial aspartate synthesis as an essential metabolic dependency for the proliferation of pancreatic tumor cells under hypoxic culture conditions. In contrast, GOT2-catalyzed aspartate synthesis is dispensable for pancreatic tumor formation *in vivo*. The dependence of pancreatic tumor cells on aspartate synthesis is bypassed in part by a hypoxia-induced potentiation of extracellular protein scavenging via macropinocytosis. This effect is mutant KRas-dependent, and is mediated by hypoxia inducible factor 1 (HIF1A) and its canonical target carbonic anhydrase-9 (CA9) through the cooption of the bicarbonate-macropinocytosis signaling axis. Our findings reveal high plasticity of aspartate metabolism and define an adaptive regulatory role for macropinocytosis by which mutant KRas tumors can overcome nutrient deprivation under hypoxic conditions.

## MAIN

Nutrient scarcity and low oxygen levels are hallmarks of the tumor microenvironment^1^. To survive under these stress conditions, cancer cells leverage oncogenic alterations that engage multiple stress-adaptive responses and facilitate the production of energy and biosynthetic precursors necessary for cell division^2^. Thus, identification of the molecular underpinnings of stress-adaptive pathways can provide insight into potential cancer cell-specific vulnerabilities.

Activating KRas mutations are among the most common oncogenic events in human cancers^3^. Through the engagement of multiple effector mechanisms, mutant KRas directs the reprogramming of cellular metabolism to favor macromolecular synthesis and enhance nutrient acquisition capabilities. Indeed, KRas activation is associated with an increase in glucose and glutamine utilization, which supports the synthesis of critical precursors for cell proliferation such as nucleotides and lipids^4,5^. Furthermore, mutant KRas cells display enhanced autophagic flux^6,7^ and macropinocytosis^8^, facilitating intracellular protein catabolism and extracellular protein degradation, respectively. These adaptive mechanisms are particularly relevant for mutant KRAS-driven pancreatic ductal adenocarcinoma (PDAC), which is severely nutrient limited owing to a dense desmoplastic stroma, and collapsed blood vessels^9^. However, their relative contribution to the fitness of pancreatic cancer cells under metabolic stress conditions is yet to be determined.

As oxygen is an essential substrate for many biosynthetic reactions^10^, hypoxia leads to multiple metabolic limitations that pancreatic cancer cells need to overcome. We have previously shown that, in tumor cell lines, hypoxia limits access to electron acceptors essential for the synthesis of aspartate and aspartate-derived nucleotides^11^. Indeed, aspartate production is limiting for a subset of hypoxic tumors and increasing its availability is sufficient to promote their growth^11,12^. In the present study, we set out to identify cellular mechanisms utilized by pancreatic tumor cells to overcome aspartate limitation under hypoxic conditions.

## RESULTS

### Metabolic routes for aspartate synthesis in hypoxic pancreatic cancer cells

PDAC often grows in a hypoxic microenvironment due to poorly functioning vasculature^13,14^. To identify metabolic genes PDAC cells depend on to proliferate under hypoxia, we performed a focused, pooled CRISPR screen of 200 rate-limiting and cancer-relevant metabolic genes in mouse PDAC cells grown under normoxic (21%) or hypoxic conditions (0.5%) (**Fig. 1a**). Among the essential genes was *HPRT*, a key enzyme in the nucleotide salvage pathway, consistent with a limiting role for nucleotides under hypoxia. The top scoring gene on the positive end of this screen was *MTHFD2* (**Fig. 1a**), consistent with its role as a major NADH source through serine catabolism when respiration is inhibited^15^. Most notably, our screen identified the mitochondrial aspartate aminotransferase (*GOT2*) as the top scoring gene necessary for PDAC cell growth under hypoxia (**Fig. 1a, Extended Data Fig. 1a**). These results are consistent with our previous findings demonstrating that human and mouse PDAC cells display substantial decreases in aspartate and nucleotide (UTP, CTP) levels when grown in low-oxygen conditions^11^ (**Extended Fig. 1b**). Confirming the screen results, GOT2 null cells (**Extended Data Fig. 1c**) proliferated substantially slower than their sgRNA-resistant GOT2 cDNA-expressing counterparts in the context of low oxygen (**Fig. 1b, Extended Data Fig. 1d**). As aspartate can be synthesized by both cytosolic (GOT1) and mitochondrial (GOT2) aspartate transaminases^16^, our results identify a preferential dependency of PDAC cells on mitochondrial aspartate synthesis under hypoxic culture conditions.

**Figure 1.**
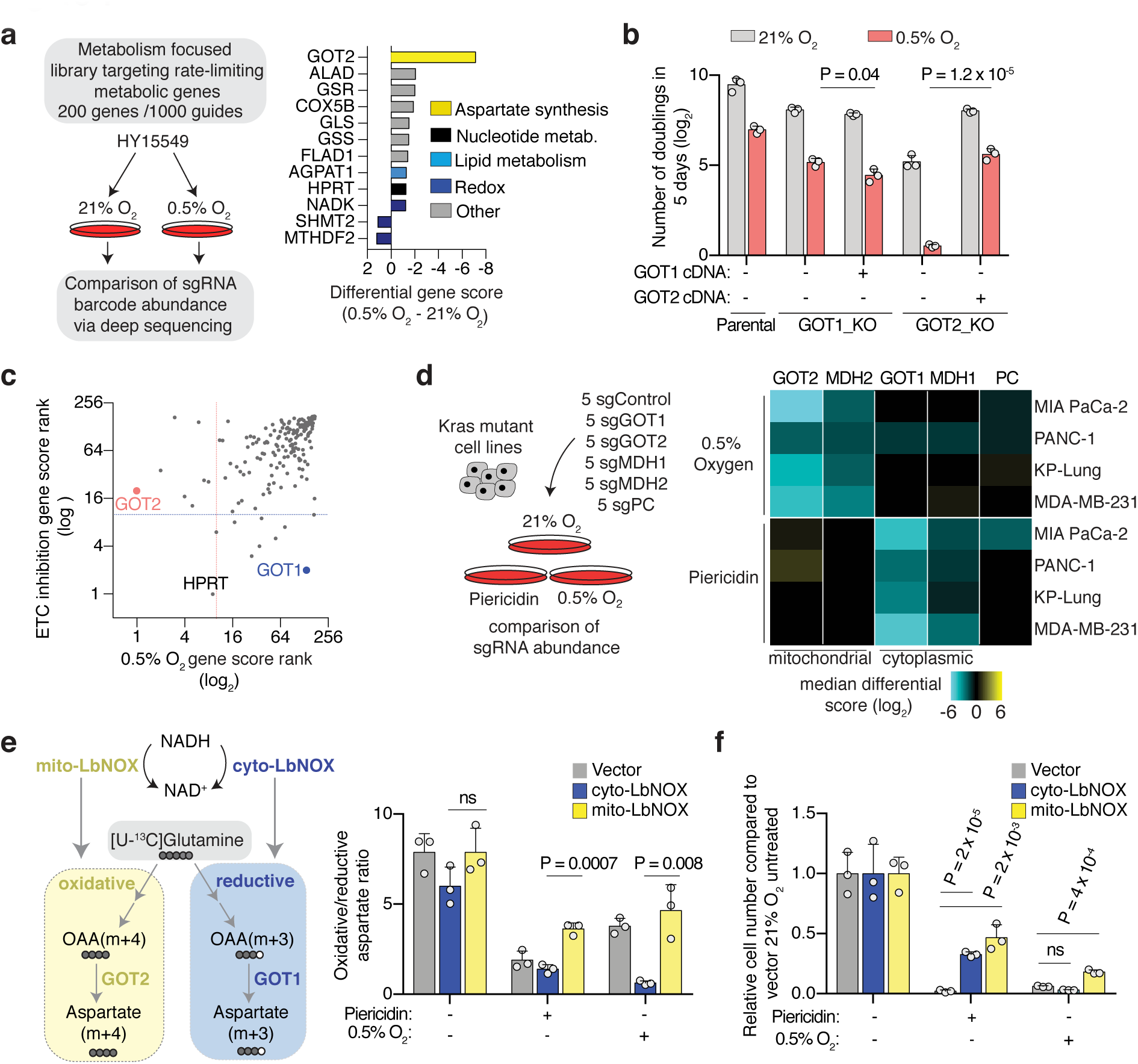
The metabolic route for aspartate synthesis in hypoxic pancreatic cancer cells. **(a)** Scheme of the focused CRISPR-Cas9 based genetic screen in a Ras mutant mouse PDAC line (HY15549) grown under normoxic (21% O_2_) or hypoxic conditions (0.5% O_2_) for 14 population doublings (left). Top scoring genes and their differential gene scores in the focused CRISPR screen (right). **(b)** Fold change in cell number (log_2_) of indicated parental, GOT1 and GOT2 knockout HY15549 cell lines expressing sgRNA-resistant GOT1 or GOT2 cDNA and grown for 5 days under normoxic (21% O_2_) or hypoxic conditions (0.5% O_2_). **(c)** Plot of gene score ranks of hypoxia and ETC inhibitor CRISPR screens in HY15549 cells. **(d)** Scheme of the *in vitro* sgRNA competition assay performed in KRas-mutant cancer cell lines transduced with a pool of 5 control sgRNAs (sgControl) and sgRNAs targeting five enzymes involved in aspartate metabolism (GOT1, GOT2, MDH1, MDH2 and PC) (left). Heat map showing median differential gene scores in the indicated cell lines upon low oxygen (0.5% O_2_) or treatment with piericidin (Pier.), an ETC inhibitor (right). **(e)** Scheme depicting the oxidative (yellow) and reductive (blue) routes from which aspartate can be synthesized from glutamine, and the activation of each pathway by mito- and cyto-LbNOX enzymes. Filled circles represent ^13^C atoms derived from [U-^13^C]-L-glutamine (left). Oxidative (m+3) to reductive (m+4) aspartate ratio in HY15549 cell lines grown under 21% O_2,_ 0.5% O_2_ or in the presence of piericidin (50 nM) (right). **(f)** Relative fold change in cell number in HY15549 cell lines transduced with a vector control, cyto-LbNOX or mito-LbNOX grown for 5 days under 0.5% O_2_ or in the presence of piericidin (50 nM). **b**, **e**, **f**, Bars represent mean ± s.d.; **b**, **d, e, f,***n* = 3 biologically independent samples. Statistical significance was determined by two-tailed unpaired *t*-test.

As oxygen is the final electron acceptor for the mitochondrial electron transport chain (ETC) and severe hypoxia may impact ETC activity^17^, we next asked whether PDAC cells rely on similar metabolic routes to grow under pharmacologic ETC inhibition. Consistent with the impairment of respiration under hypoxic conditions, treating cells with ETC inhibitors (**Extended Data Fig. 1e**) and culturing them under hypoxia trigger comparable metabolic signatures in PDAC cell lines (**Extended Data Fig. 1f, g**). However, in contrast to the similarities in metabolomic profiles, many genetic essentialities were unique to each condition (**Fig. 1c, Extended Data Fig. 1h**). In particular, while mitochondrial GOT2 is essential for PDAC cell growth under hypoxic conditions, PDAC cells depend on the cytoplasmic GOT1 under ETC inhibition^18^ (**Extended Data Fig. 1i**). Confirming the screen results, loss of GOT1, but not GOT2, strongly sensitizes cells to ETC inhibition in individual proliferation assays (**Extended Data Fig. 1j-l**). These effects are independent of the role of GOTs in redox homeostasis, as increasing aspartate availability by expressing an intracellular guinea pig asparaginase (gpASNase)^12^ is sufficient to rescue the proliferation defects upon hypoxia or ETC inhibition in both parental and GOT1/2 knock out cells (**Extended Data Fig. 1m, n**). To determine whether this is a generalizable phenomenon, we performed a small-scale sgRNA competition assay in multiple PDAC cell lines with sgRNAs targeting control genomic regions within each of the 4 genes in the aspartate malate shuttle (*GOT1*, *MDH1*, *GOT2*, *MDH2*), or pyruvate carboxylase (*PC*)^19^ (**Fig. 1d**). These competition assays revealed that knocking-out mitochondrial enzymes of the aspartate-malate shuttle (GOT2 and MDH2) strongly inhibits the growth of these cell lines under hypoxia, whereas loss of the cytoplasmic enzymes (GOT1 and MDH1) sensitizes them to ETC inhibition, suggesting that distinct compartmentalized routes operate for aspartate synthesis under ETC inhibition versus hypoxia (**Fig. 1d**).

### Mitochondrial NAD^+^ is limiting for PDAC cell proliferation under hypoxia

Impairment of respiration limits NAD^+^ for tricarboxylic acid (TCA) cycle progression, thereby hindering production of oxaloacetate, the precursor for aspartate synthesis^18,20^. As PDAC cells depend on the mitochondrial, but not cytoplasmic, aspartate transaminase under hypoxia, we hypothesized that mitochondrial NAD^+^ may be more limiting for PDAC cell proliferation under hypoxic conditions. To test this idea, we exploited mitochondria- or cytoplasmic-localized versions of the water-forming NADH oxidase LbNOX, a bacterial enzyme that recycles NADH into NAD^+^ (**Fig. 1e, Extended Data Fig. 2a**)^21^. Consistent with previous reports^21^, expression of mito-LbNOX regenerates mitochondrial NAD^+^ and allows cells to maintain TCA cycle activity, supporting oxidative glutamine metabolism in spite of ETC inhibition or low oxygen (**Fig. 1e, Extended Data Fig. 2b**). Conversely, cytoplasmic-LbNOX (cyto-LbNOX) expression regenerates cytoplasmic NAD^+^ and promotes reductive metabolism of glutamine (**Fig.1e, Extended Data Fig. 2b**). Using this system, we found that only mito-LbNOX expression promotes cell proliferation under hypoxia (**Fig. 1f, Extended Data Fig. 2c**) and this effect requires the presence of GOT2 (**Extended Data Fig. 2d**). Accordingly, in the absence of GOT2, increasing aspartate availability by gpASNase expression rescued hypoxia-induced proliferation arrest (**Extended Data Fig. 2d**). These results suggest that the selective dependence of PDAC cells on GOT2 for proliferation under hypoxia might be imposed by the limiting levels of mitochondrial NAD^+^ under these conditions. Of note, the anti-proliferative effects of pharmacologic ETC inhibition were rescued by both cyto- or mito-LbNOX (**Fig. 1f, Extended Data Fig. 2c**) further indicating that ETC inhibition and hypoxia confer distinct metabolic dependencies.

### Aspartate synthesis by GOTs is redundant for PDAC tumor formation

While cancer cells in culture rely on GOT1 and GOT2 under ETC inhibition or hypoxia, respectively, the relative contributions of these enzymes to tumor growth *in vivo* are not clear. To address this, we generated tumors from isogenic GOT1 and GOT2 knock out PDAC cell lines (murine HY15549^22^ and human MIA PaCa-2) and their counterparts expressing sgRNA-resistant GOT1/2 cDNA. Loss of GOT2 did not decrease the size of MIA PaCa-2 tumors and only modestly impacted those derived from HY15549 cancer cell line (**Fig. 2a**). Similarly, GOT1 knock out tumors did not show any significant change in tumor growth when compared to GOT1 cDNA-expressing counterparts (**Fig. 2a**). *In vivo* sgRNA competition assays confirmed that the lack of essentiality of GOT1 or GOT2 can be generalized to other PDAC cell lines in tumors implanted subcutaneously or orthotopically (**Extended Data Fig. 3a**) as well as a patient-derived xenograft (PDX) (**Extended Data Fig. 3b**). Building upon these results, we first investigated the possible contribution of GOT1 to aspartate synthesis in the absence of GOT2 (**Fig. 2b**). GOT1-mediated aspartate synthesis is only possible when cells have an exogenous NAD^+^ source^18,20^. Consistent with this requirement, supplementation of culture media with pyruvate, an exogenous electron acceptor that can regenerate NAD^+^, rescued the anti-proliferative effects of hypoxia in GOT2-KO cells (**Extended Data Fig. 3c**) and in parental cells in a manner dependent on GOT1 activity (**Extended Data Fig. 3d**). Notably, pyruvate is one of the metabolites secreted by human pancreatic stromal cells (hPSC)^23,24^. Indeed, addition of hPSC-conditioned media to HY15549 cells promotes hypoxic cell proliferation similar to pyruvate supplementation (**Extended Data Fig. 3e**). We next sought to determine whether pyruvate uptake might enable the growth of GOT2-deficient PDAC tumors *in vivo*. Cancer cells take up pyruvate through monocarboxylate transporters, in particular MCT1^25^ (**Fig. 2b**). Pharmacologic inhibition of MCT1 (**Extended Data Fig. 3e**) or its genetic loss in GOT2 knock out PDAC cells (**Extended Data Fig. 3f**) halted hypoxic proliferation under physiological pyruvate levels^26^ (**Fig. 2c**). Consistent with the significant contribution of extracellular pyruvate to PDAC tumor growth, MCT1/GOT2 double KO cells form significantly smaller tumors compared to those expressing MCT1 cDNA (**Fig. 2d**). These results suggest that environmental pyruvate may provide sufficient NAD^+^ to support aspartate synthesis and tumor growth.

**Figure 2.**
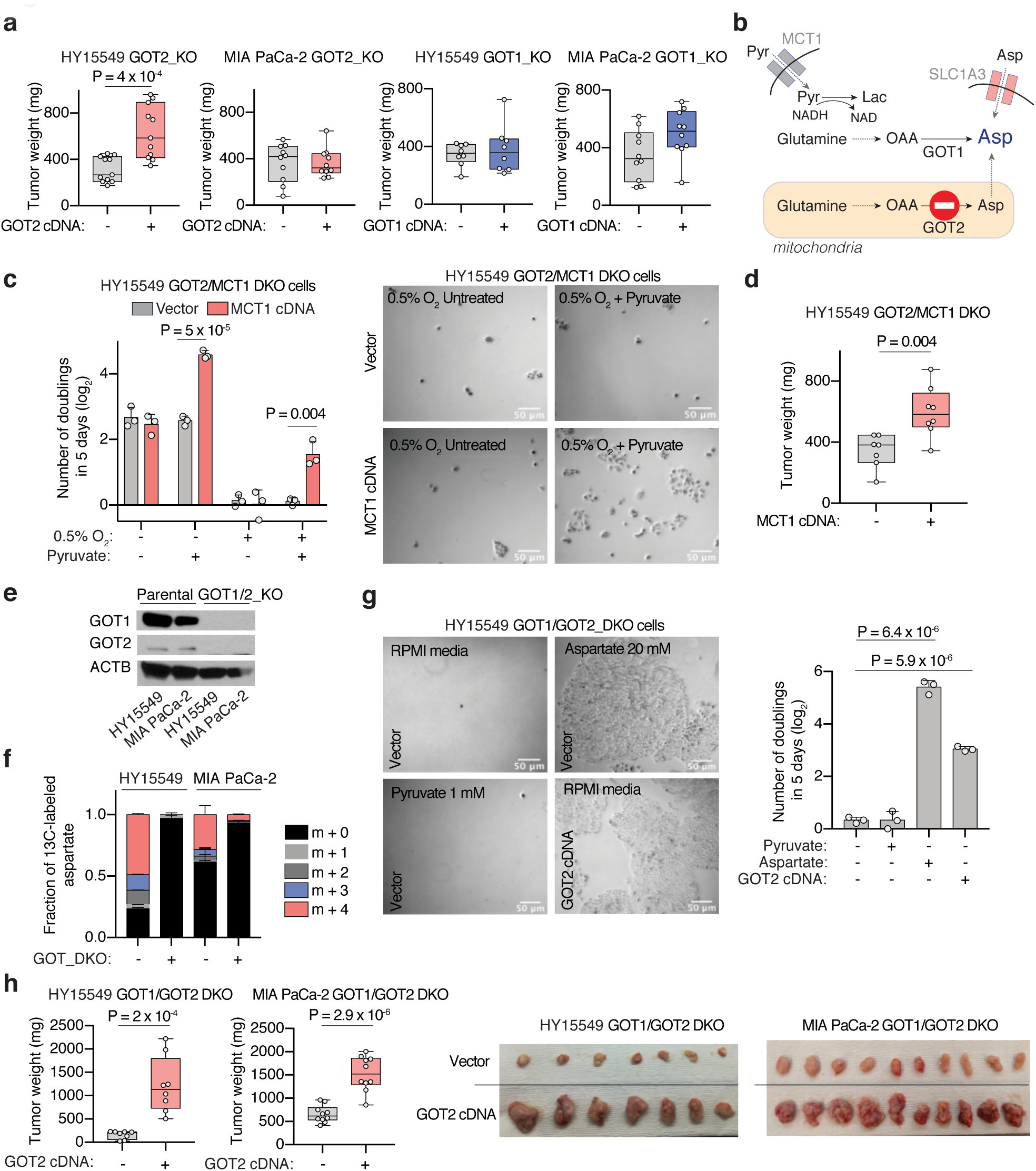
The plasticity of aspartate metabolism in PDAC tumors. **(a)** Weights of subcutaneous tumor xenografts derived from the indicated GOT1- and GOT2-knockout cell lines transduced with a control vector or an sgRNA-resistant GOT1/2 cDNA. **(b)** Scheme depicting metabolic pathways to compensate for the loss of aspartate synthesis in GOT2-knockout cells. Pyruvate can be taken up by monocarboxylate transporter 1 (MCT1, grey), and in turn acts as an exogenous electron acceptor, enabling GOT1-mediated aspartate synthesis in the cytosol. Alternatively, environmental aspartate could be obtained by cells expressing a plasma membrane aspartate transporter (SLC1A3, pink). **(c)** Fold change (log_2_) in cell number of GOT2/MCT1 double-knockout HY15549 cells transduced with a control vector or an sgRNA-resistant MCT1 cDNA after 5 days of growth in the absence or presence of pyruvate (100 μM) (left). Representative bright-field micrographs of indicated cells under 0.5% O_2_ in the presence or absence of 100 μM pyruvate. Scale bar = 50 μm (right). **(d)** Weights of subcutaneous tumor xenografts derived from the GOT2/MCT1 double-knockout HY15549 cells transduced with a control vector or an sgRNA-resistant MCT1 cDNA. **(e)** Immunoblot analysis of GOT1 and GOT2 in parental or GOT1/GOT2 double knockout HY15549 and MIA PaCa-2 cells. ACTB was used as loading control. **(f)** Fraction of ^13^C-labeled aspartate in parental and GOT1/GOT2 double knockout HY15549 and MIA PaCa-2 cells cultured for 8 hours in the presence of 0.5 mM [U-^13^C]-L-glutamine. **(g)** Representative bright-field micrographs of GOT1/GOT2 double knockout HY15549 cells transduced with a control vector or an sgRNA-resistant GOT2 cDNA grown for 5 days in indicated culture conditions. Scale bar = 50 μm (left). Fold change (log_2_) in cell number of GOT1/GOT2 double knockout HY15549 cells transduced with a control vector or an sgRNA-resistant GOT2 cDNA grown for 5 days in the absence and presence of pyruvate (1 mM) or aspartate (20 mM). **(h)** Weights of subcutaneous tumor xenografts derived from the indicated GOT1/GOT2 double knockout cell lines transduced with a control vector or sgRNA-resistant GOT2 cDNA (left). Representative images of indicated tumors (right). **a, d, h,** boxes represent the median, and the first and third quartiles, and the whiskers represent the minima and maxima of all data points. **c**, **f**, **g,** Bars represent mean ± s.d. **a, d, h,** *n* = 7–12 biologically independent samples. **c**, **f**, **g,** *n* = 3 biologically independent samples. Statistical significance was determined by two-tailed unpaired *t*-test.

To assess directly whether de novo aspartate synthesis is redundant for tumor formation, we generated HY15549 and MIA PaCa-2 PDAC cell lines lacking both aspartate aminotransferases (GOT1/2 DKO), thus auxotrophic for aspartate (**Fig. 2e**). Glutamine labeling confirmed that these GOT1/2 DKO cells do not synthesize aspartate de novo (**Fig. 2f**). Consequently, these cells can only be grown in media supplemented with supraphysiological levels of aspartate (20 mM), highlighting their inability to grow *in vitro* (**Fig. 2g, Extended Data Fig. 3g**). While aspartate synthesis by GOTs is necessary for survival in culture, GOT1/2 DKO PDAC cells still form tumors, albeit smaller (**Fig. 2h**), indicating the presence of alternative sources for aspartate acquisition *in vivo*. Of note, aspartate levels in MIA PaCa-2 DKO tumors were comparable to those of cells expressing GOT2 cDNA (**Extended Data Fig. 3h**). These results suggest that tumor aspartate availability is maintained by both synthesis and alternative extracellular sources *in vivo*.

### Macropinocytosis provides sufficient aspartate to enable PDAC cell growth under hypoxic culture conditions

To identify potential extracellular sources of aspartate we first considered the possibility that the loss of aspartate biosynthesis could be compensated by the uptake of extracellular aspartate via plasma membrane aspartate/glutamate SLC1A family transporters^11^ (**Fig. 2b**). However, none of the major SLC1A aspartate transporters are expressed in PDAC cell lines (**Extended Data Fig. 3i, j**)^27^. An alternative mechanism that may enable cells to obtain aspartate from extracellular sources is the scavenging of albumin via macropinocytosis, a mutant KRas dependent process^8^. Indeed, the macropinocytic uptake of albumin, the most abundant protein in interstitial fluids, and its subsequent lysosomal degradation has been shown to generate free amino acids^8,28–30^. We therefore tested the effects of bovine serum albumin (BSA) supplementation on aspartate levels and growth of PDAC cells. The growth defect of KRas mutant PDAC cell lines under hypoxic conditions was partly rescued by BSA supplementation (**Fig. 3a**). Metabolomic analysis of MIA PaCa-2 cells grown under hypoxia in the presence or absence of BSA identified aspartate as the only amino acid whose levels significantly decreased in response to hypoxia, and the addition of BSA was sufficient to restore its levels (**Fig. 3b**). In addition, BSA dramatically increased the levels of pyrimidine synthesis intermediates (dihydroorotate, orotate and uridine-monophosphate, UMP) (**Fig. 3c**), for which aspartate is a precursor^11^. These results suggest that macropinocytosis could provide sufficient aspartate to promote proliferation of KRAS-mutant cancer cells under hypoxic conditions (**Extended Data Fig. 4a**).

**Figure 3.**
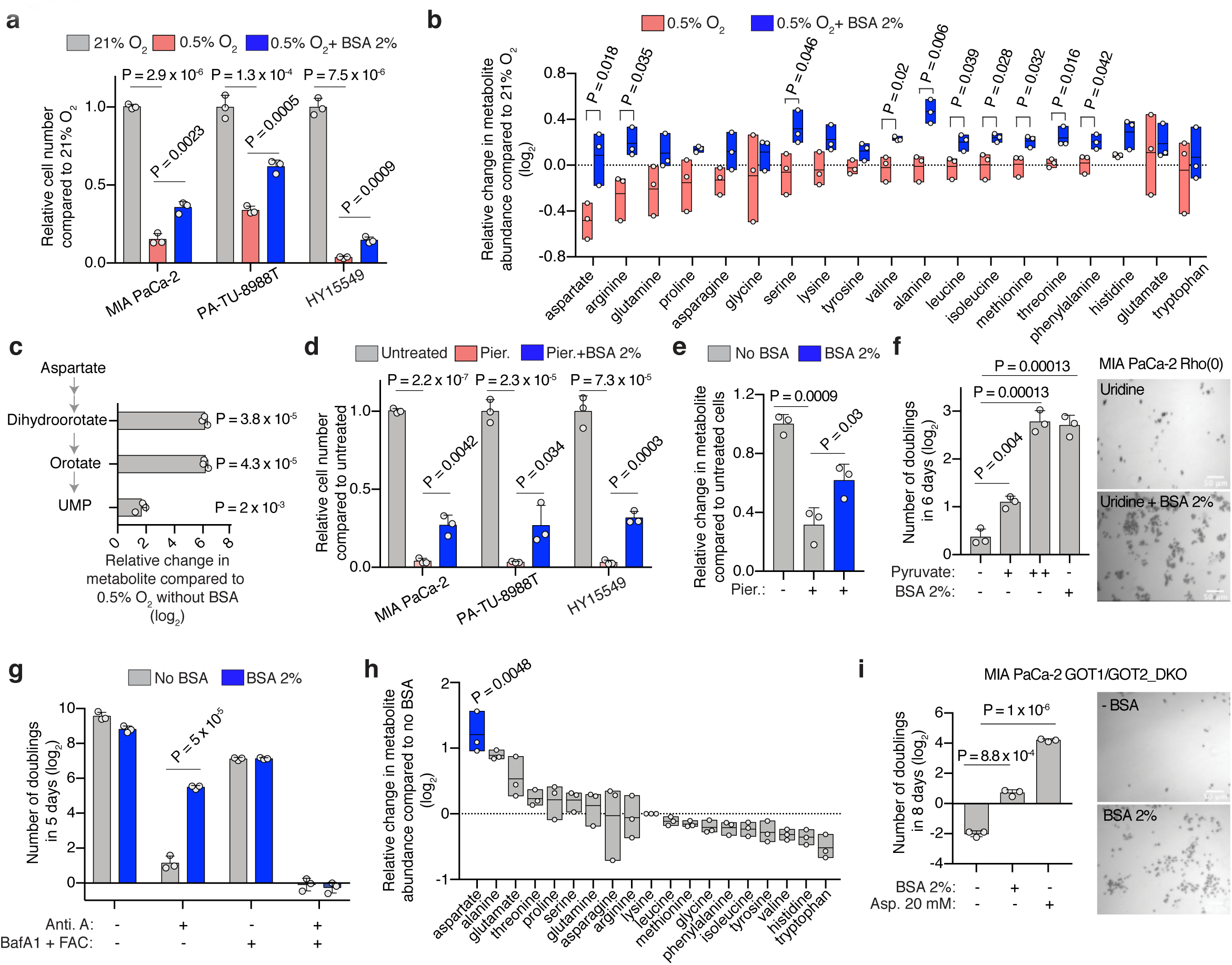
Macropinocytosis provides sufficient aspartate to enable PDAC cell growth under hypoxic culture conditions. **(a)** Relative fold change in cell number of indicated PDAC cell lines grown for 5 days under hypoxic conditions (0.5%) in the presence or absence of 2% BSA to those cultured under normoxia (21%). **(b)** Relative change (log_2_) in amino acid abundance of MIA PaCa-2 cells grown under 0.5% O_2_ in the presence (blue) or absence (pink) of 2% BSA to those cultured under normoxia (21%). **(c)** Relative change (log_2_) in the indicated pyrimidine synthesis metabolic intermediates in MIA PaCa-2 cells grown under 0.5% O_2_ with 2% BSA to those without BSA treatment. **(d)** Relative fold change in cell number of indicated PDAC cell lines treated with piericidin (Pier.; MIA PaCa-2 100 nM; PA-TU-8998T 100 nM; HY15549 30 nM) for 5 days in the presence or absence of 2% BSA to corresponding untreated cells. **(e)** Relative aspartate abundance of MIA PaCa-2 cells treated with piericidin (150 nM) in the presence and absence of 2% BSA to that of untreated cells. Metabolite abundance is normalized by total protein levels. **(f)** Fold change in cell number (log_2_) of MIA PaCa-2 Rho(0) cells grown for 6 days in media supplemented with uridine (50 μg/mL) in the presence or absence of pyruvate (+, 100 μM; ++, 1 mM) or 2% BSA (left). Representative bright-field micrographs of MIA PaCa-2 Rho(0) cells in the presence or absence of 2% BSA. Scale bar = 50 μm (right). **(g)** Fold change in cell number (log_2_) of HY15549 cells grown for 5 days in the absence and presence of 2% BSA, antimycin A (Anti. A.; 100 nM) or Bafilomycin A1 (BafA1; 10 nM + ferric ammonium citrate (FAC) 0.1 μg/mL). **(h)** Relative change (log_2_) in amino acid abundance of GOT1/GOT2 double knockout MIA PaCa-2 cells grown for 48 hrs with 2% BSA to cells cultured in the absence of BSA. Aspartate is highlighted in blue. **(i)** Fold change in cell number (log_2_) of GOT1/GOT2 double knockout MIA PaCa-2 cells grown for 8 days in regular RPMI media and media supplemented with 2% BSA or aspartate (20 mM) (left). Representative bright-field micrographs of GOT1/GOT2 double knockout MIA PaCa-2 cells in the presence or absence of BSA 2%. Scale bar = 50 μm (right). **a**, **c**, **d**, **e, f, g, i,** Bars represent mean ± s.d.; **b, h,** boxes represent the median, and the first and third quartiles. **a-i,** *n* = 3 biologically independent samples. Statistical significance was determined by two-tailed unpaired *t*-test.

We next sought to determine whether albumin-mediated rescue of cell growth is restricted to hypoxia or can be generalized to other conditions where aspartate levels are limiting. We therefore tested the ability of albumin to rescue the anti-proliferative effects of ETC inhibition. Similar to hypoxia, BSA supplementation partly overcame the anti-proliferative effects of a complex I inhibitor (**Fig. 3d**) and the drop in aspartate levels (**Fig. 3e**) in PDAC cell lines. Likewise, the addition of BSA to MIA PaCa-2 cells that are ETC-defective by virtue of lacking mtDNA^31^ (MIA PaCa-2 ρ^0^) (**Extended Data Fig. 4b**) enabled their proliferation to the same extent observed upon pyruvate supplementation (**Fig. 3f**). Consistent with BSA exerting its effects through macropinocytic uptake and lysosomal degradation, treatment of HY-15549 with Bafilomycin A1 (BafA1), an inhibitor of lysosomal acidification^32^, prevents the proliferation rescue by BSA under ETC inhibition (**Fig. 3g, Extended Data Fig. 4c**). Of note, in line with previous findings, these experiments require iron supplementation to maintain viability of cells without lysosomal acidity^33^. Finally, consistent with albumin as a sufficient source for aspartate, BSA supplementation increases aspartate levels (**Fig. 3h**) and enables proliferation (**Fig. 3i**) of MIA PaCa-2 GOT1/2 DKOs cells that are auxotrophic for aspartate. This effect is dependent on the presence of mutant Ras, as addition of BSA to GOT1/2 DKO cells generated from BXPC-3 cell line with wild type (WT) Ras alleles (**Extended Data Fig. 4d**), allows cell proliferation only upon expression of oncogenic KRas (**Extended Data Fig. 4e**). Collectively, these experiments suggest that protein scavenging via macropinocytosis contributes sufficient aspartate for growth when aspartate is limiting.

### Hypoxia upregulates macropinocytosis in mutant KRAS cells *in vivo* and *in vitro*

Given our findings linking macropinocytosis to an increase in aspartate levels under hypoxic conditions, we set out to investigate the relationship between hypoxia and macropinocytosis. To this end, tumor xenografts derived from MIA PaCa-2 cells were co-stained with tetramethylrhodamine-labeled high molecular weight dextran (TMR-dextran), a marker of macropinocytosis; and pimonidazole, a non-toxic 2-nitroimidazole hypoxia marker^34^ (**Fig. 4a**). A marked increase in TMR-dextran uptake by tumor cells, identified by positive staining for cytokeratin 8 (CK8), was observed in areas of high pimonidazole staining, suggesting that tumor hypoxia correlates with elevated macropinocytosis (**Fig. 4b, c**). Consistent with these *in vivo* data, hypoxia also caused a pronounced increase in the uptake of TMR-dextran in KRas-mutant PDAC cell lines *in vitro* (**Fig. 4d, e**). The hypoxia-dependent increase in macropinocytic uptake was restricted to KRas-mutant cells, and the expression of oncogenic KRas in WT Ras cells was sufficient to confer this dependency, implicating a direct role for mutant KRAS in linking hypoxia to the modulation of macropinocytosis levels (**Fig. 4f-i, Extended Data Fig. 5a**). Moreover, in KRas mutant cells incubated with DQ-BSA, the hypoxia-induced macropinocytic uptake was accompanied by an increase in DQ-BSA fluorescence, indicating a corresponding increase the proteolytic degradation of BSA (**Fig. 4j, k, Extended Data Fig. 5b-e**). Thus, the upregulation of macropinocytosis under hypoxic conditions might serve as an adaptive response to nutrient deprivation imposed by low oxygen conditions.

**Figure 4.**
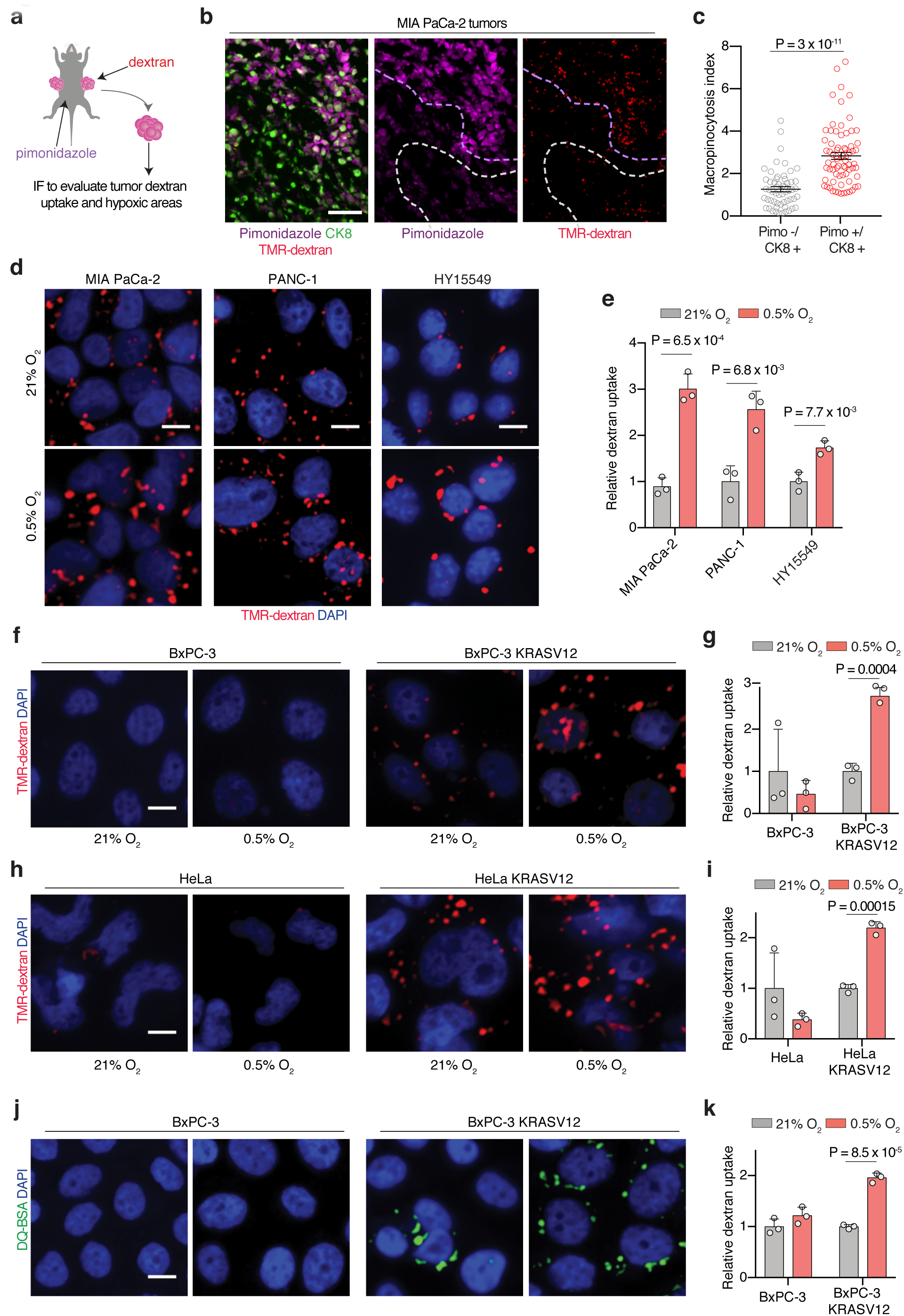
Hypoxia upregulates macropinocytosis in mutant KRAS cells *in vivo* and *in vitro*. **(a)** Schematic of *in vivo* assay for administering intratumoral TMR-dextran and intraperitoneal pimonidazole injection to identify macropinosomes and hypoxic regions of tumors, respectively. **(b)** Hypoxic tumor areas display more macropinocytosis than neighboring non-hypoxic areas. Representative images from sections of MIA PaCa-2 xenograft tumor tissue showing macropinosomes labeled with TMR-dextran (red), tumor cells immunostained with anti-CK8 (green), and pimonidazole detected with anti-pimonidazole (purple). Dashed lines indicate high (purple) and low (white) areas of pimonidazole staining. Scale bar = 50 μm. **(c)** Quantification of macropinocytic uptake in **(b)** showing tumor macropinocytosis index in hypoxic tumor areas (CK8+/Pimo+, red) relative to non-hypoxic tumor areas (CK8+/Pimo-, grey). **(d, e)** Representative images (**d**) and quantification (**e**) of TMR-dextran (red) uptake in MIA PaCa-2, PANC-1 and HY15549 cells under normoxia (21% O_2_) and hypoxia (0.5% O_2_). Nuclei are labeled with DAPI (blue). Scale bar, 10 μm. In **e**, data are presented relative to normoxia of the respective cell line. **(f-i)** Hypoxia-induced macropinocytosis depends on oncogenic RAS. Representative images (**f, h**) and quantification (**g, i**) of TMR-dextran (red) uptake in BxPC-3 cells (**f, g**) and HeLa cells (**h, i**) without and with expression of doxycycline-inducible KRasG12V grown in normoxia (21% O_2_) or hypoxia (0.5% O_2_). Nuclei are labeled with DAPI (blue). Scale bar, 10 μm. In **g**, **i**, data are presented relative to normoxia of the respective cell line. **(j-k)** Representative images (**j**) and quantification (**k**) of DQ-BSA fluorescence (green) in BxPC-3 cells without and with expression of doxycycline-inducible KRasG12V grown in normoxia (21% O_2_) or hypoxia (0.5% O_2_). In **k**, data are presented relative to normoxia of the respective cell line. **c, e, g, i, k,** Bars represent mean ± s.e.m. At least 300 (**c**) or 500 (**e, g, i, k**) cells were quantified in each biological replicate (*n* = 3). Statistical significance was determined by two-tailed unpaired *t*-test.

### Hypoxia-inducible factor 1A (HIF1A) is necessary and sufficient for hypoxia-dependent upregulation of macropinocytosis

HIF1 is a heterodimeric, basic helix-loop-helix transcription factor that is composed of HIF1A, the alpha subunit, and the aryl hydrocarbon receptor nuclear translocator (ARNT), the beta subunit^35^. HIF1A is considered the master transcriptional regulator of cellular responses to hypoxia^36^, raising the possibility that it might play a role in the potentiation of macropinocytosis under hypoxic conditions. Indeed, KRas mutant cells rendered deficient for HIF1A or ARNT by CRISPR-Cas9 gene editing (**Fig. 5a, b, Extended Data Fig. 6a**) failed to display an increase in macropinocytosis in response to hypoxia (**Fig. 5c, d, Extended Data Fig. 6b, c**). Conversely, mutant KRas cells transduced with a constitutively active HIF1A variant (HIF1A P402A/P564A)^37^ (**Fig. 5e, Extended Data Fig. 6d**) displayed an enhancement of macropinocytosis under standard atmospheric conditions, and a robust potentiation of the increase in macropinocytosis triggered by low oxygen (**Fig. 5f, g, Extended Data Fig. 6e, f**). Consistent with these results, stabilization of alpha subunits of HIFs by chemical inhibition of prolyl-hydroxylases (PHDs)^38^ also induces TMR-dextran uptake (**Extended Data Fig. 6g-i**) only in PDAC cells with oncogenic Ras mutations (**Extended Data Fig. 6i**). Next, we tested the effects of loss- and gain-of function of HIF1A on macropinocytosis *in vivo* using tumor xenografts established from HIF1A/ARNT knockout cells and HIF1A P402/P564A expressing cells. In agreement with the *in vitro* data, loss of HIF1A/ARNT abrogated the potentiation of macropinocytosis in hypoxic tumor regions (**Fig. 5h, Extended Data Fig. 6j**), whereas constitutive activation of HIF1A resulted in the enhancement of macropinocytosis in both hypoxic and non-hypoxic regions (**Figure 5i, j**). Collectively, these results indicate that the HIF1A/ARNT axis is a necessary and sufficient mediator of hypoxia-induced potentiation of macropinocytosis.

**Figure 5.**
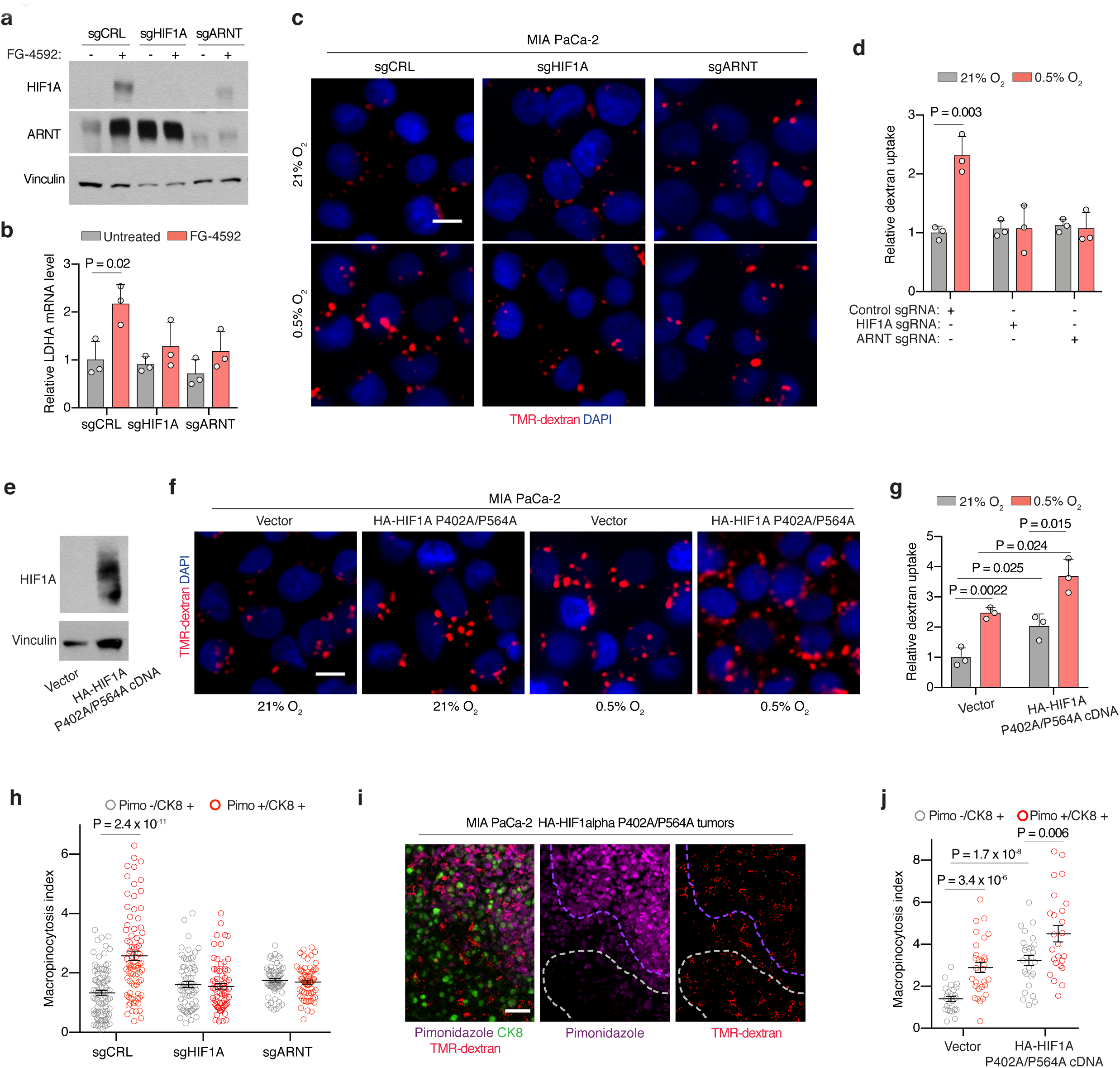
Hypoxia-inducible factor 1A (HIF1A) is necessary and sufficient for hypoxia-dependent upregulation of macropinocytosis. **(a)** Immunoblot analysis of HIF1A and ARNT proteins in the indicated MIA PaCa-2 HIF1A and ARNT knockout cell lines treated with the prolyl-hydroxylase (PHD)-inhibitor FG-4592 (100 μM, 72 hrs) as shown. Vinculin was used as a loading control. FG-4592 is used to stabilize HIFs and confirm pathway disruption. **(b)** Relative mRNA levels of HIF1A-target lactate dehydrogenase A (LDHA) mRNA expression in the indicated MIA PaCa-2 knockout cells treated with FG-4592 (100 μM for 72 hrs) compared to untreated cells. **(c, d)** Representative images (**c**) and quantification (**d**) of TMR-dextran (red) uptake in the indicated MIA PaCa-2 knockout cell lines under normoxia (21% O) or hypoxia (0.5% O_2_). Nuclei are labeled with DAPI (blue). Scale bar, 10 μm. In **d**, data are presented relative to values obtained for normoxic control cells. **(e)** Immunoblot analysis of HIF1A in MIA PaCa-2 cells expressing a constitutively stable HIF1A isoform (HA-HIF1A P402/P564A) cDNA. Vinculin was used as a loading control. **(f, g)** Representative images (**f**) and quantification (**g**) of TMR-dextran (red) uptake in MIA PaCa-2 cells transduced with vector control or HA-HIF1A P402/P564A cDNA under normoxia (21% O_2_) or hypoxia (0.5% O_2_). In **g**, data are presented relative to normoxia cells transduced with a control vector. **(h)** Quantification of macropinocytic uptake from sections of xenograft tumors derived from MIA PaCa-2 cells transduced with a control sgRNA or sgRNAs targeting HIF1A or ARNT showing tumor macropinocytic index in hypoxic tumor areas (CK8+/Pimo+, red) and non-hypoxic tumor areas (CK8+/Pimo-, grey). Data is presented relative to non-hypoxic areas in tumors arising from cells transduced with a control sgRNA. **(i)** Representative images from sections of xenograft tumors derived from MIA PaCa-2 cells expressing HA-HIF1A P402/P564A cDNA showing macropinosomes labeled with TMR-dextran (red), tumor cells immunostained with anti-CK8 (green), and pimonidazole detected with anti-pimonidazole (purple). Nuclei are labeled with DAPI (blue). Dashed lines indicate high (purple) and low (white) areas of pimonidazole staining. Scale bar = 50 μm. **(j)** Quantification of macropinocytic uptake in **(i)** showing tumor macropinocytic index in hypoxic tumor areas (CK8+/Pimo+, red) and non-hypoxic tumor areas (CK8+/Pimo-, grey). Data is presented relative to non-hypoxic areas in tumors derived from cells transduced with a control vector. **b, d, g,** Bars represent mean ± s.e.m.; **h, j**, At least 500 (**c, d, g**) or 300 (**h, j**) cells were quantified in each biological replicate (*n* = 3). Statistical significance was determined by two-tailed unpaired *t*-test.

### The HIF1A target CA9 is required for the upregulation of macropinocytosis in response to hypoxia

We next sought to determine the mechanism underlying the requirement of HIF1A for hypoxia-induced upregulation of macropinocytosis. Among the canonical HIF1A downstream targets is carbonic anhydrase IX (CA9), a transmembrane zinc metalloenzyme functioning as the catalyst of reversible hydration of carbon dioxide to bicarbonate ions and protons^39,40^. Indeed, we observed that CA9 is upregulated in hypoxic regions of mutant KRas tumors (**Extended Data Fig. 7a, b**) and in mutant KRas cell lines expressing constitutively stable HIF1A (**Extended Data Fig. 7c**). We have recently reported that oncogenic Ras-induced stimulation of macropinocytosis is dependent on the engagement of bicarbonate signaling axis that is initiated by an increase in the expression of the electroneutral bicarbonate transporter SLC4A7^41^. The increase in bicarbonate influx that ensues leads to the activation of bicarbonate-dependent soluble adenylate cyclase (sAC) and its downstream effector protein kinase A (PKA)^42^, which in turn leads to a series of alterations in membrane dynamics that are required for macropinocytosis^41^. Building upon these findings, we hypothesized that CA9, by virtue of its capacity to increase bicarbonate availability, could play a critical role in coupling hypoxia to the upregulation of macropinocytosis. To test this hypothesis, we generated CA9-deficient mutant KRas cells using CRISPR-Cas9 gene editing (**Extended Data Fig. 7d**) and assessed their macropinocytic activity *in vitro* and *in vivo*. In cells cultured under atmospheric oxygen conditions, loss of CA9 had no impact on macropinocytosis levels. However, the potentiation of macropinocytosis observed under hypoxic conditions was abrogated in CA9-deficient cells (**Fig. 6a, b, Extended Data Fig. 7e, f**). The same CA9 dependence was observed *in vivo*, wherein the upregulation of macropinocytosis characteristic of hypoxic regions of the tumor failed to take place in tumor xenografts derived from CA9-deficient cells (**Fig. 6c, d**). Lastly, overexpression of CA9 in KRas mutant cell lines (**Extended Data Fig. 7g**) resulted in significant stimulation of baseline macropinocytosis activity, and further potentiated it in response to hypoxia (**Fig. 6e, f, Extended Data Fig. 7h, i**). Together, these results indicate that HIF1A-dependent induction of CA9 expression is an essential step in the adaptation process that enables mutant KRas cells to upregulate their macropinocytic activity under hypoxic conditions. The relevance of the catalytic product of CA9, bicarbonate, to this process is supported by the observation that siRNA-mediated knockdown of the bicarbonate transporter SLC4A7 or treatment of cells with a pan-bicarbonate transporter inhibitor, S0895, resulted in a partial or complete blockade of macropinocytosis under hypoxia, respectively (**Fig. 6g; Extended Data Fig. 7j**). Consistent with the engagement of this bicarbonate signaling axis by way up CA9 upregulation, treatment of cells with a PKA inhibitor, H89, resulted in the loss of the potentiating effect of hypoxia on macropinocytosis (**Extended Data Fig. 7k**).

**Figure 6.**
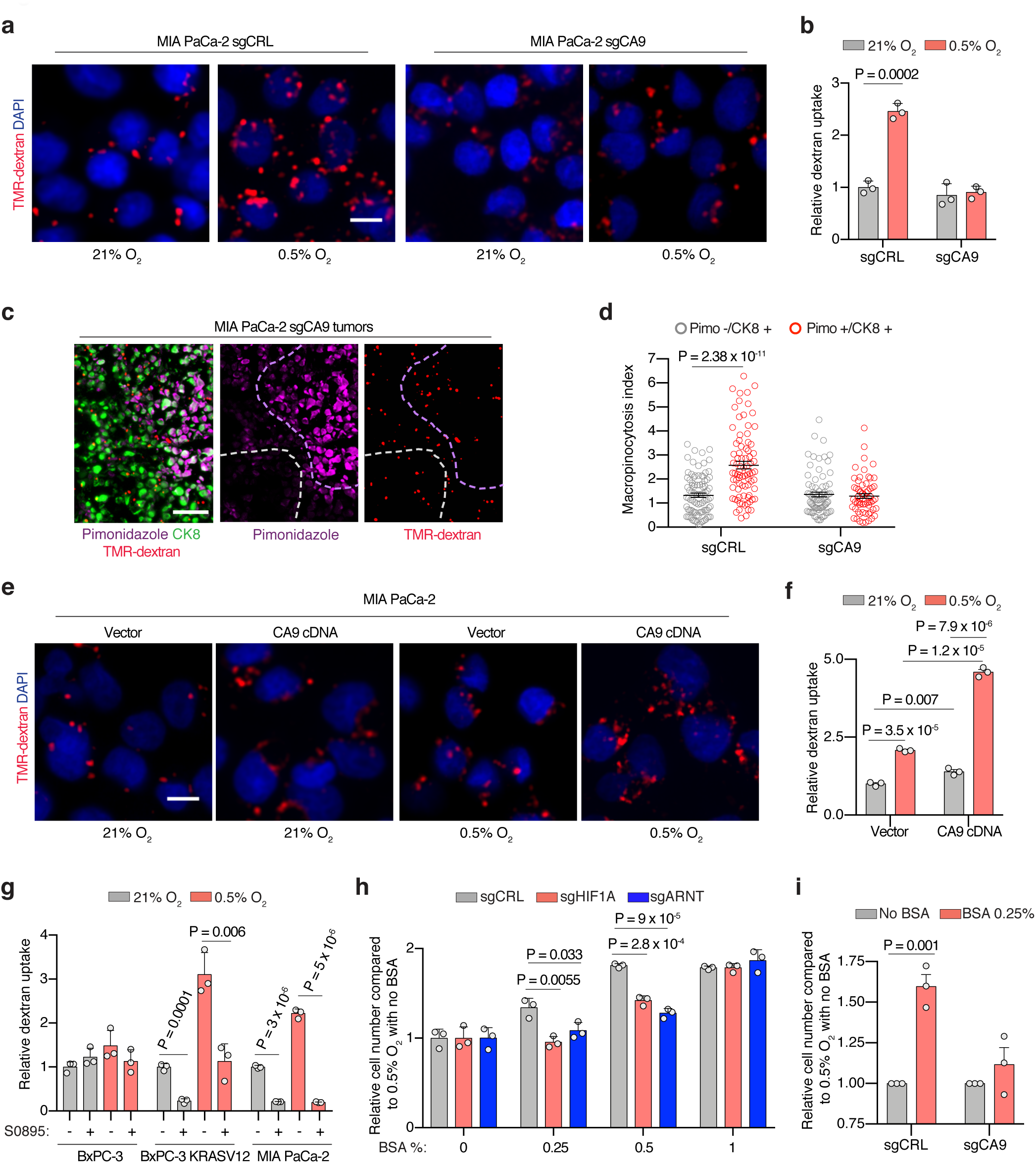
The HIF1A target CA9 is required for the upregulation of macropinocytosis in response to hypoxia. **(a, b)** Representative images (**a**) and quantification (**b**) of TMR-dextran (red) uptake in control sgRNA or sgCA9-transduced MIA PaCa-2 cells under normoxia (21% O_2_) or hypoxia (0.5% O_2_). Nuclei are labeled with DAPI (blue). Scale bar, 10 μm. Data are presented relative to values obtained for normoxic control cells. **(c)** Representative images from sections of xenograft tumors derived from MIA PaCa-2 cells transduced with an sgRNA targeting CA9 showing macropinosomes labeled with TMR-dextran (red), tumor cells immunostained with anti-CK8 (green), and pimonidazole detected with anti-pimonidazole (purple). Nuclei are labeled with DAPI (blue). Dashed lines indicate high (purple) and low (white) areas of pimonidazole staining. Scale bar = 50 μm. **(d)** Quantification of macropinocytic uptake in (**c**) showing tumor macropinocytic index in hypoxic tumor areas (CK8+/Pimo+, red) and non-hypoxic tumor areas (CK8+/Pimo-, grey). Data is presented relative to that of non-hypoxic areas in tumors arising from cells transduced with control sgRNA. **(e, f)** Representative images (**e**) and quantification (**f**) of TMR-dextran (red) uptake in control or CA9 overexpressing MIA PaCa-2 cells under normoxia (21% O_2_) or hypoxia (0.5% O_2_). Nuclei are labeled with DAPI (blue). Scale bar, 10 μm. In **f**, data are presented relative to values obtained for normoxic control cells. **(g)** Quantification of macropinocytic uptake in the indicated cell lines treated with the bicarbonate transporter inhibitor S0895 (15 μM) as shown under normoxia (21% O_2_) or hypoxia (0.5% O_2_). Data are presented relative to normoxia of the respective untreated cell line. **(h)** Fold change in cell number of the indicated MIA PaCa-2 cell lines grown for 5 days under hypoxia (0.5% O_2_) with increasing concentrations of BSA (0%-1%). Data are presented relative to hypoxic cells transduced with a control sgRNA and cultured without BSA. **(i)** Fold change in cell proliferation in the indicated MIA PaCa-2 cell lines after 5 days of proliferation under hypoxia (0.5% O_2_) and in the absence or presence of BSA (0.25%). Data are presented relative to values obtained for hypoxic cells cultured without BSA. **b, f, g,** Bars represent mean ± s.e.m.; At least 500 (**b, d, g**) or 300 (**f**) cells were quantified in each biological replicate (*n* = 3). **h-i,** Bars represent mean ± s.d.; **h-i,** *n* = 3 biologically independent samples. Statistical significance was determined by two-tailed unpaired *t*-test.

### Potentiation of macropinocytosis by HIF1A/CA9 axis supports cell growth under nutrient limitation

We have previously shown that the deleterious effect of amino acid deprivation on the growth of mutant KRas cells can be rescued by supplementing the growth medium with albumin^8,43^. Furthermore, we and others have demonstrated that albumin must be supplied at near physiological levels (1-3%) to support the survival and growth of mutant KRas cells under these conditions. Since the rescuing capacity of albumin in these settings is dependent on its macropinocytic uptake, we reasoned that hypoxia-induced potentiation of macropinocytosis could reduce the threshold levels of albumin required for cell growth. To test this idea, we assessed the dependence of cell growth on albumin levels under hypoxic conditions. Mutant KRas cells displayed a significant growth advantage when subjected to hypoxia in the presence of sub-physiological levels of albumin (**Fig. 6h, i**). Furthermore, this adaptive response was dependent on the HIF1A/CA9 axis, and by extension, on the potentiation of macropinocytosis by hypoxia, as it could not be mounted by cells deficient for HIF1A, ARNT, and CA9 (**Fig. 6h, i**). Altogether, these experiments highlight the functional relevance of the HIF1A/CA9-driven stimulated macropinocytosis in PDAC cell proliferation under nutrient limitation.

## CONCLUSION

As an essential substrate for more than 140 metabolic reactions, oxygen serves many critical anabolic and catabolic functions in mammalian cells. To overcome metabolite limitations imposed by hypoxia, most cancer cells engage stress-adaptive responses and utilize alternative nutrient sources. In this study we explored the nature of these adaptive responses in the context of aspartate availability. While the limiting role of aspartate under both hypoxic and respiration impaired conditions has been established^11,18,20^, our results reveal that cancer cells utilize distinct subcellular routes of aspartate synthesis to accommodate each stress. PDAC cells depend on mitochondrial aspartate synthesis via GOT2 under hypoxia, whereas rely on cytoplasmic aspartate production in response to pharmacologic ETC inhibition. This is likely in part due to differences in compartmentalized NAD^+^ limitation under these distinct conditions, but also may be mediated by hypoxia-specific downstream effectors. The use of compartment-specific pathways as an adaptation response to nutrient deprivation has previously been described for one carbon metabolism^44^, and may represent a general strategy exploited by cancer cells to enhance their metabolic adaptive capabilities.

As the limiting role of aspartate has been universally observed in culture, aspartate transaminases have previously been suggested as potential therapeutic targets^5^. However, our findings demonstrate that *in vivo* PDAC cells can partly overcome the complete loss of de novo aspartate production. This apparent discrepancy can be attributed to both cell extrinsic and cell intrinsic factors: neighboring cells secrete pyruvate, thereby providing reducing equivalents to cancer cells; and cancer cells themselves upregulate macropinocytosis to improve aspartate acquisition. Given the presence of multiple pathways to address aspartate limitation, the relative contribution of each to tumor growth needs to be carefully assessed before considering this pathway for anti-cancer therapies.

While this study focuses on the acquisition of aspartate through macropinocytosis under limiting oxygen conditions, macropinocytosis itself is a non-selective bulk-fluid uptake process, and therefore likely would enhance the capacity of hypoxic tumor cells to import other extracellular solutes and macromolecules. Consistent with this notion, previous work has indicated that lipid scavenging is stimulated by hypoxic culture conditions and can be phenocopied in normoxic cells by Ras mutation^45^. Thus, increased macropinocytosis in the context of Ras mutation and its exacerbation by hypoxia would be predicted to enhance lipid scavenging, fueling membrane replication and cell division. In addition, a low oxygen environment causes necrosis leading to the accumulation of necrotic cellular debris that can enter the cells via macropinocytosis^46^ and provide a robust source of important bioenergetic substrates. Lastly, the adaptive response of cells to hypoxia involves inhibition of mammalian target of rapamycin complex 1 (mTORC1)^47^. In oncogenic Ras transformed cells, inhibition of mTORC1 has been shown to increase the endolysosomal-mediated catabolism of proteins that enter the cells via macropinocytosis^30^. Collectively, hypoxic reprogramming of nutrient scavenging via macropinocytosis emerges as a versatile adaptive mechanism by which mutant Ras tumors could improve metabolic fitness.

We have previously shown that bicarbonate availability couples oncogenic Ras signaling to the molecular machinery that controls macropinocytosis^41^. The present study identifies this coupling mechanism as the point of intersection between hypoxia and macropinocytosis through the upregulation of CA9 and the ensuing increase in bicarbonate production. Significantly, high levels of CA9 expression in PDAC tumors have been shown to be associated with reduced survival^48^. Moreover, both small molecules and antibodies targeting CA9 are currently at various stages of clinical development^49^. Therefore, the link between CA9 expression and nutrient uptake may represent a new targetable vulnerability of mutant Ras tumors, and suggests that inhibition of this enzyme in conjunction with other anabolic pathways could yield effective treatments for PDAC.

## ACKNOWLEDGEMENTS

We thank all members of the Bar-Sagi and Birsoy labs for helpful suggestions. The KP mouse cancer cell lines are gifts from N. Bardeesy and T. Papagiannakopoulos. We would like to thank the New York University Langone Medical Center (NYULMC) Experimental Pathology Research Laboratory, which is partially supported by NIH/NCI 5370 P30CA16087.This research is supported by an NIH K99/R00 to J.G.-B. (1K99CA248838-01). K.B. was supported by NCI (DP2 OD024174-01), AACR NextGen Grant, Pershing Square Sohn foundation and the Breast Cancer Research Foundation; and is a Searle, Pew-Stewart and Sidney Kimmel Scholar. This work was supported by NIH/NCI Grant CA210263 (D.B.-S).

## AUTHOR CONTRIBUTIONS

D.B.-S., S.P., K.B. and J.G.-B. conceived the project and designed the experiments. J.G.-B. and S.P. performed most of the experiments with assistance of L.B., Y.L. and M.S. K.L. performed computational analysis. H.A, B.R. and H.M. performed analysis of metabolite profiling experiments. R.T.W. performed real time PCR experiments. R.F.H. provided hPSCs. E.S. provided PDAC PDXs and N.Y. performed experiments with them. D.B.-S., K.B. and J.G.-B wrote and edited the manuscript with help from M.A.B. and L.T.

## EXTENDED DATA FIGURE LEGENDS

**Extended Data Figure 1.**
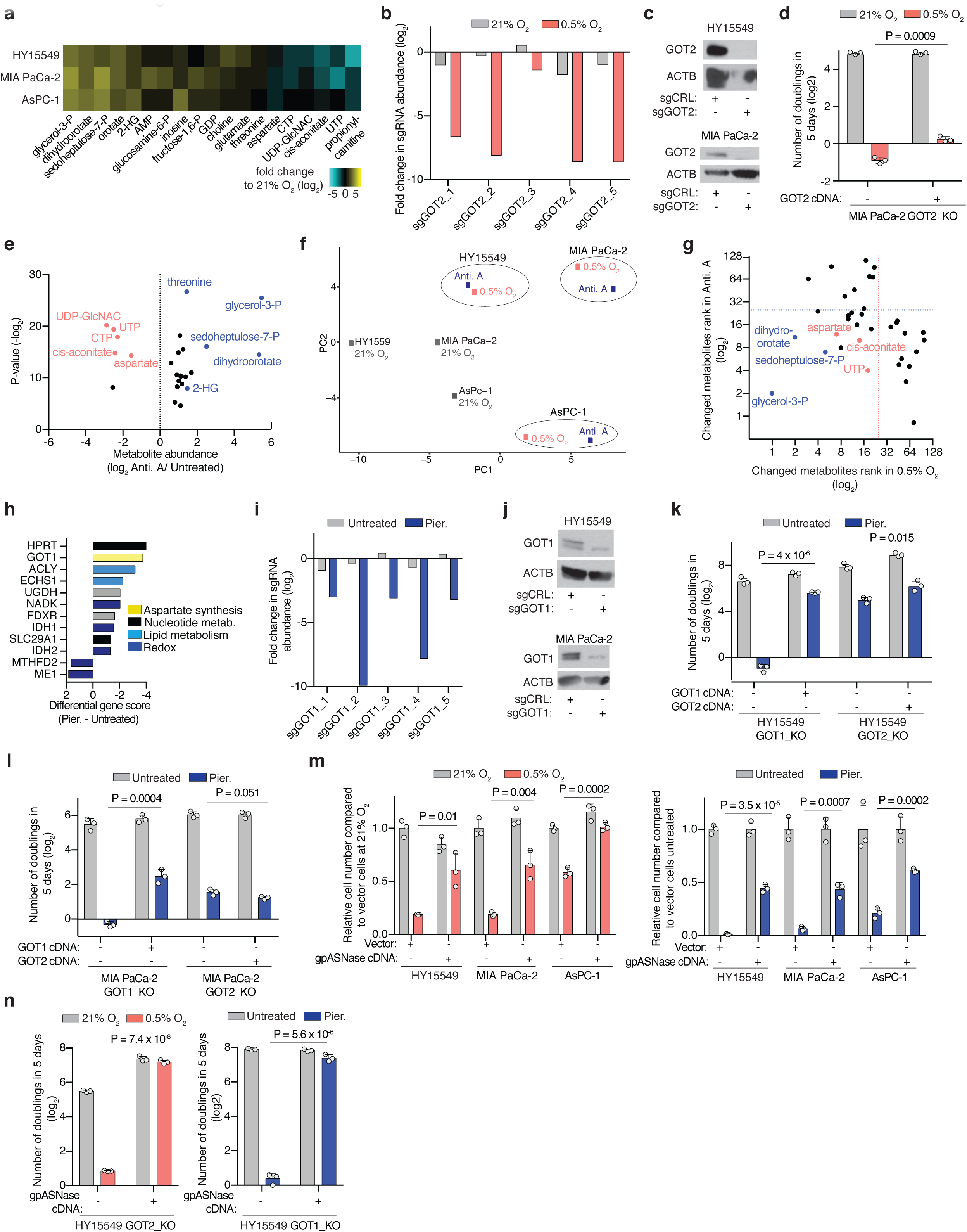
Hypoxia and ETC inhibition trigger similar metabolic signatures in PDAC cells. **(a)** Individual sgRNA scores for GOT2 under normoxia (21% O_2_) and hypoxia (0.5% O_2_). **(b)** Heat map showing fold changes (log2) of polar metabolites significantly changed across three PDAC cell lines exposed to low oxygen (0.5% O_2_) compared to metabolite levels of cells grown at 21% O_2_. **(c)** Immunoblot analysis of GOT2 in HY15549 and MIA PaCa-2 GOT2 knockout cell lines compared to parental cells transduced with a control sgRNA. ACTB was used as a loading control. **(d)** Fold change in cell number (log_2_) of indicated MIA PaCa-2 cell line expressing a control vector or an sgRNA-resistant GOT2 cDNA grown under indicated conditions for 5 days. **(e)** Common metabolite changes across three PDAC cell lines upon ETC inhibition. Data is shown as the ratio of metabolite abundance in antimycin A-treated cells to untreated controls (log_2_ x-axis) versus significance of difference between the median abundances of each metabolite in both groups (-log_2_p-value, y-axis). **(f)** Principal component analysis (PCA) reveals that metabolic changes in PDAC cells exposed to 0.5% O_2_ (pink) or antimycin A treatment (blue) are similar. **(g)** Ranks of metabolites most significantly changed (p value < 0.01, dotted lines) across 3 PDAC cell lines under ETC inhibition (upper left quartile) or 0.5% O_2_ (lower right quartile). Metabolites significantly accumulated (blue) or depleted (pink) under both conditions are in the lower left quartile (right). **(h)** Top scoring genes and their differential gene scores in HY15549 cells treated with piericidin (Pier.) compared to untreated cells. **(i)** Individual sgRNA scores for GOT1 in HY15549 cells in the presence or absence of piericidin. **(j)** Immunoblot analysis of GOT1 in HY15549 and MIA PaCa-2 cell lines expressing GOT1 sgRNA compared to those with a control sgRNA. ACTB was used as a loading control. **(k)** Fold change in cell number (log_2_) of indicated HY15549 cell lines expressing a control vector or an sgRNA-resistant GOT1 or GOT2 cDNA grown for 5 days in the presence or absence of piericidin (Pier., 20 nM). **(l)** Fold change in cell number (log_2_) of indicated MIA PaCa-2 cell lines expressing a control vector or an sgRNA-resistant GOT1 or GOT2 cDNA grown for 5 days in the presence or absence of piericidin (Pier., 20 nM). **(m)** Relative fold change in cell number of indicated PDAC cell lines grown for 5 days under 0.5% O_2_ (left) or in the absence and presence of piericidin (Pier., 20-100 nM) (right) to untreated cells cultured at 21% O_2_. **(n)** Fold change in cell number (log_2_) of indicated HY15549 cell lines transduced with a control vector or gpASNase cDNA grown for 5 days under the indicated oxygen levels (left) or in the absence and presence of piericidin (Pier. 50 nM) (right). **b, i,** bars represent individual sgRNA scores; **d**, **k**, **l, m, n,** Bars represent mean ± s.d‥ **d**, **k**, **l, m, n,** *n* = 3 biologically independent samples. Statistical significance was determined by two-tailed unpaired *t*-test.

**Extended Data Figure 2.**
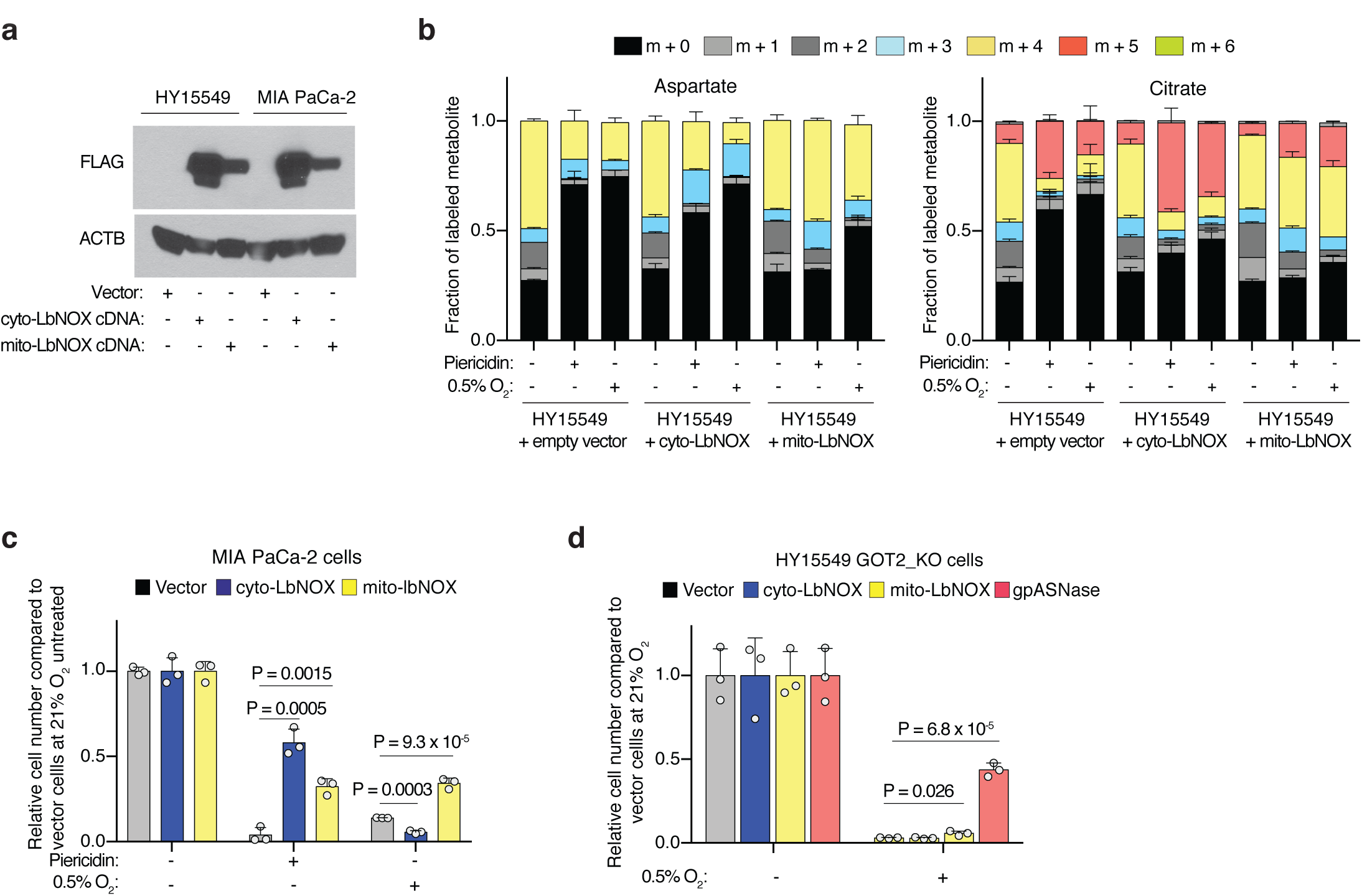
Mitochondrial NAD^+^ is limiting for PDAC cell proliferation under hypoxia. **(a)** Immunoblot analysis of FLAG-tagged cytoplasmic (cyto-) and mitochondrial (mito-) LbNOX heterologous expression in the indicated cell lines. ACTB was used as a loading control. **(b)** Fraction of labelled aspartate (left) and citrate (right) derived from labelled glutamine in control and cyto- or mito-LbNOX expressing HY15549 cells cultured for 8 h with [U-^13^C]-Glutamine (500 μM) with piericidin treatment (50 nM) or under 0.5% oxygen. Colors indicate mass isotopomers. **(c)** Expression of LbNOX enzymes rescues proliferation of MIA PaCa-2 cells under piericidin treatment (50 nM), whereas only mito-LbNOX expression rescues proliferation under 0.5% O_2_. Data is shown as fold change in cell number in the indicated cell lines grown for 5 days relative to untreated cells cultured at 21% O_2_. **(d)** Relative fold change in cell number of HY15549 GOT2-knockout cells transduced with a control vector or the indicated cDNAs grown for 5 days under hypoxia (0.5% O_2_) to those cultured under normoxia (21% O_2_). **b-d,** Bars represent mean ± s.d. **b-d,** *n* = 3 biologically independent samples. Statistical significance was determined by two-tailed unpaired *t*-test.

**Extended Data Figure 3.**
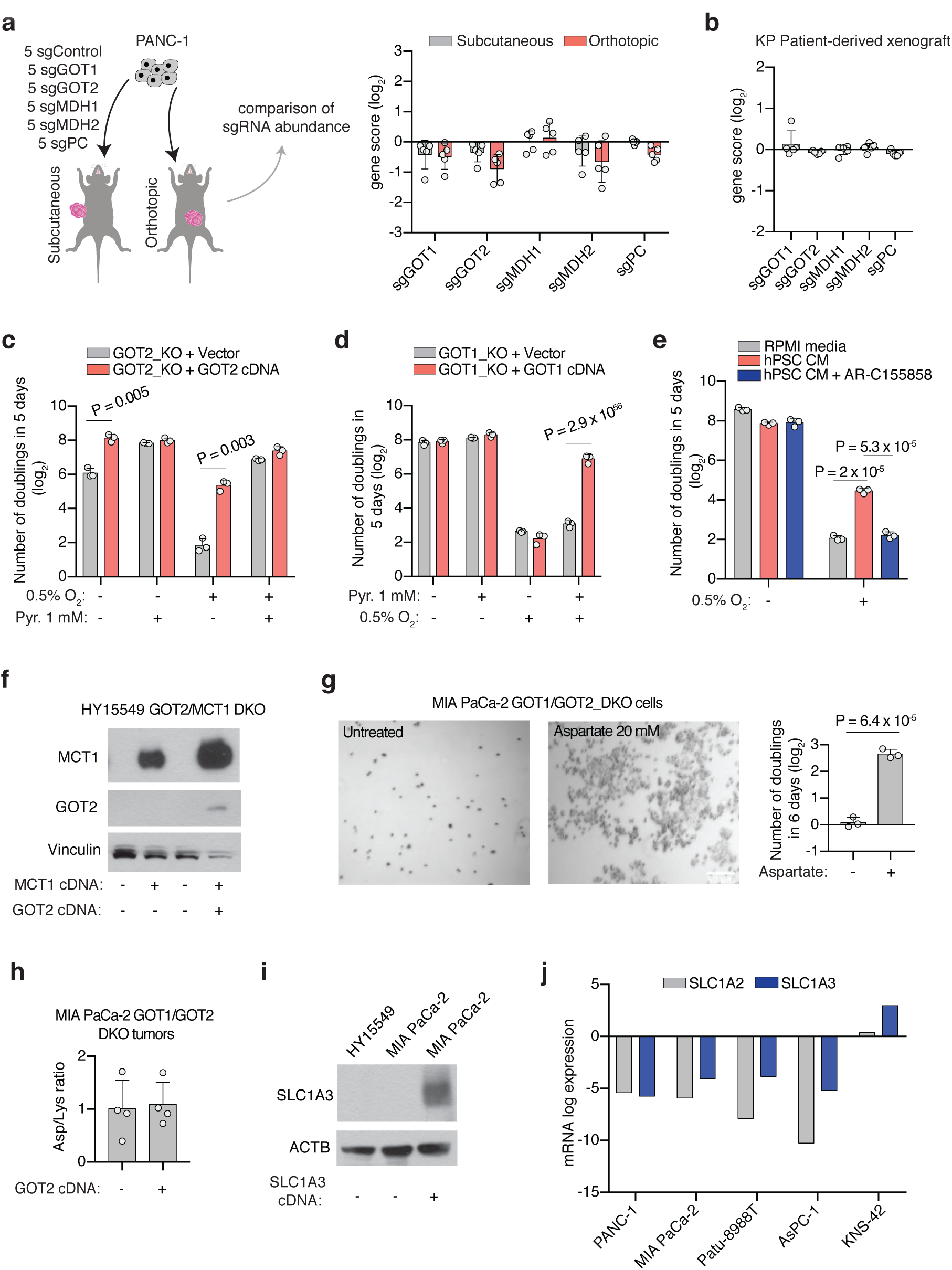
Aspartate synthesis by GOTs is essential in culture but redundant for *in vivo* tumor growth. **(a)** Scheme of the *in vivo* sgRNA competition assay performed in PANC-1 cells transduced with a pool of 5 control sgRNAs (sgControl) and sgRNAs targeting five enzymes involved in aspartate metabolism (GOT1, GOT2, MDH1, MDH2 and PC) prior to subcutaneous or orthotopic implantation (left). Bar graphs show the gene scores in subcutaneous or orthotopic PANC-1 tumors (right). Each dot represents an sgRNA targeting the indicated gene (n = 3 orthotopic tumors, 8 subcutaneous tumors). **(b)** *In vivo* sgRNA competition assay performed in a KRas-mutant PDAC patient-derived xenograft (PDX) model. Cells were transduced with a pool of 5 control sgRNAs (sgControl) and sgRNAs targeting five enzymes involved in aspartate metabolism (GOT1, GOT2, MDH1, MDH2 and PC) prior to subcutaneous implantation. Gene scores are shown, each dot represents an sgRNA targeting the indicated gene (n = 4 subcutaneous tumors). **(c)** Fold change in cell number (log_2_) of HY15549 GOT2-knockout cells transduced with a control vector or an sgRNA resistant GOT2 cDNA grown for 5 days in the absence or presence of pyruvate (Pyr., 1 mM) under hypoxic or normoxic conditions. **(d)** Fold change (log_2_) in cell number of HY15549 GOT1-KO cells transduced with a control vector or an sgRNA resistant GOT1 cDNA grown for 5 days in the absence or presence of pyruvate (Pyr., 1 mM) under hypoxic or normoxic conditions. **(e)** Fold change (log_2_) in cell number of HY15549 cells grown for 5 days in regular media or conditioned media from hPSCs (hPSC CM) in the presence or absence of the monocarboxylate transporter inhibitor AR-C155858 (5 μM). **(f)** Immunoblot analysis of MCT1 and GOT2 in HY15549 GOT2/MCT1 double knockout cells transduced with a control vector or sgRNA-resistant GOT2 or MCT1 cDNAs. Vinculin was used as a loading control. **(g)**. Representative bright-field micrographs of MIA PaCa-2 GOT1/GOT2 double knockout cells grown for 6 days in the indicated media conditions. Scale bar = 50 μm (left). Fold change in cell number (log2) of GOT1/GOT2 double knockout cells grown in the presence or absence of aspartate (20 mM). **(h)** Relative aspartate abundance of established xenografts derived from MIA PaCa-2 GOT1/GOT2 double knockout cells transduced with a control vector or GOT2 cDNA. Aspartate abundance was normalized by lysine levels. **(i)** Immunoblot shows the absence of expression of the aspartate transporter SLC1A3 in parental PDAC cell lines. Protein extract from MIA PaCa-2 cells transduced with SLC1A3 cDNA was included as a positive control. ACTB was used as a loading control. **(j)** SLC1A2 and SLC1A3 mRNA expression data from the Cancer Cell Line Encyclopedia (CCLE) (log transformed) in KP PDAC cell lines and a glioma cell line (KNS-42). **a**, **b**, **e, f, g, i, j,** Bars represent mean ± s.d. **a**, **b**, **e, f, g, i,** *n* = 3 biologically independent samples; **j,** *n* = 4 biologically independent samples. Statistical significance was determined by two-tailed unpaired *t*-test.

**Extended Data Figure 4.**
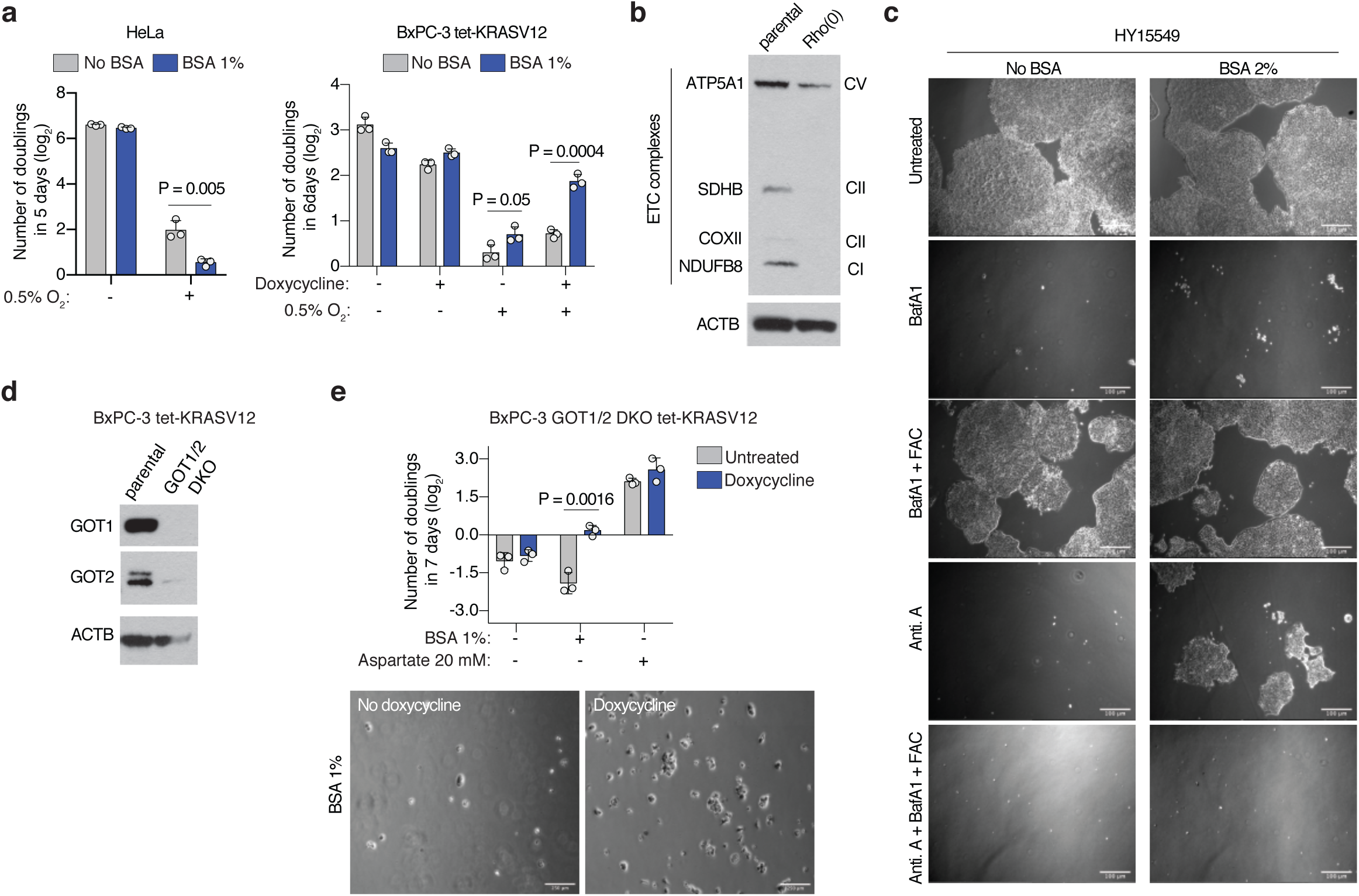
Macropinocytosis enables proliferation of KRAS-mutant cells under conditions where aspartate is limited. **(a)** Fold change in cell number (log_2_) of HeLa (left) or BxPC-3 tet-KRASV12 (right) cells cultured under indicated concentrations of oxygen for 5-6 days in the presence or absence of 1% BSA. BxPC-3 tet-KRASV12 cells were cultured in the presence or absence of doxycycline (0.1μg/mL) to activate KRASV12 expression. **(b)** Immunoblot analysis of several members of the ETC complex in parental MIA PaCa-2 cells or Rho(0) counterparts. ACTB was used as a loading control. **(c)** Representative bright-field micrographs of HY15549 cells cultured for 5 days in media with or without 2% BSA. Where indicated, cells were treated with bafilomycin A1 (BafA1, 10 nM), ferric ammonium citrate (FAC, 0.1 μg/mL) and complex III inhibitor antimycin A (Anti. A, 100 nM). Scale bar = 50 μm. **(d)** Immunoblot analysis of GOT1 and GOT2 in GOT1/2-double knockout BxPC-3 tet-KRASV12 cells compared to parental controls. ACTB was used as a loading control. **(e)** Fold change in cell number (log_2_) of GOT1/2-double knockout BxPC-3 tet-KRASV12 cells cultured in the indicated media conditions for 7 days in the presence or absence of doxycycline (0.1 μg/mL) (top). Representative bright-field micrographs of GOT1/2-double knockout BxPC-3 tet-KRASV12 cells in media supplemented with 1% BSA in the presence and absence of doxycycline (0.1 μg/mL) (bottom). **a, e,** Bars represent mean ± s.d. **a**, **e,** *n* = 3 biologically independent samples. Statistical significance was determined by two-tailed unpaired *t*-test.

**Extended Data Figure 5.**
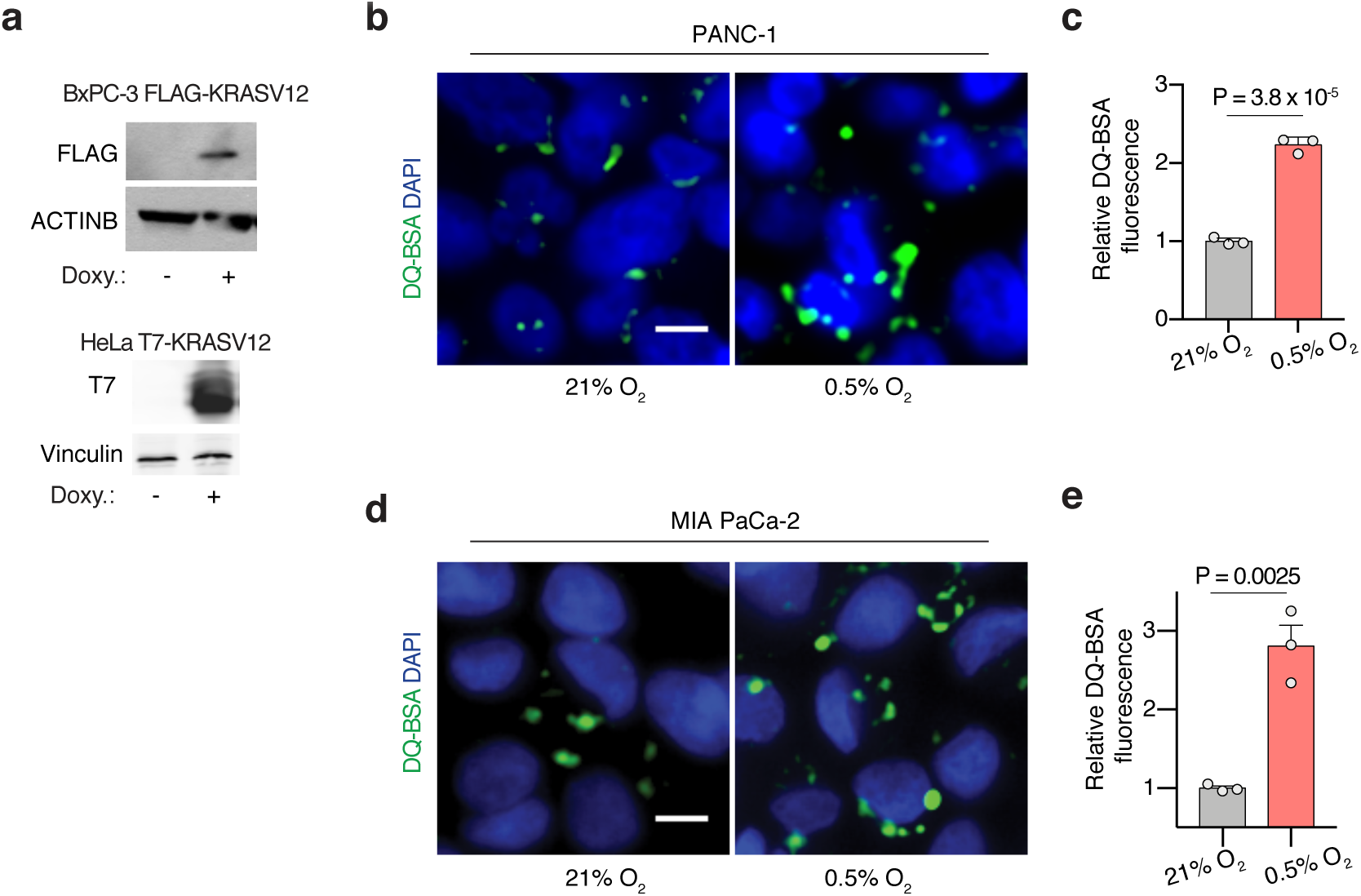
Hypoxia-induced macropinocytosis depends on oncogenic KRAS in PDACs. **(a)** Immunoblot analysis of FLAG and T7, showing inducible expression of stably transduced FLAG-tagged KRasG12V BxPC-3 cells and T7-tagged KRasG12V HeLa cells after addition of doxycycline (1 μg/mL for 2 days). ACTB or vinculin were used as a loading control. **(b-e)** Representative images (**b, d**) and quantification (**c, e**) of DQ-BSA fluorescence (green) in PANC-1 (**b, c**) and MIA PaCa-2 (**d, e**) cells under normoxia (21% O_2_) and hypoxia (0.5% O_2_). Nuclei are labeled with DAPI (blue). Scale bar, 10 μm. In **c**, **e**, data are presented relative to normoxia. **c, e,** Bars represent mean ± s.e.m. At least 500 (**c, d**) cells were quantified in each biological replicate (*n* = 3). Statistical significance was determined by two-tailed unpaired *t*-test.

**Extended Data Figure 6.**
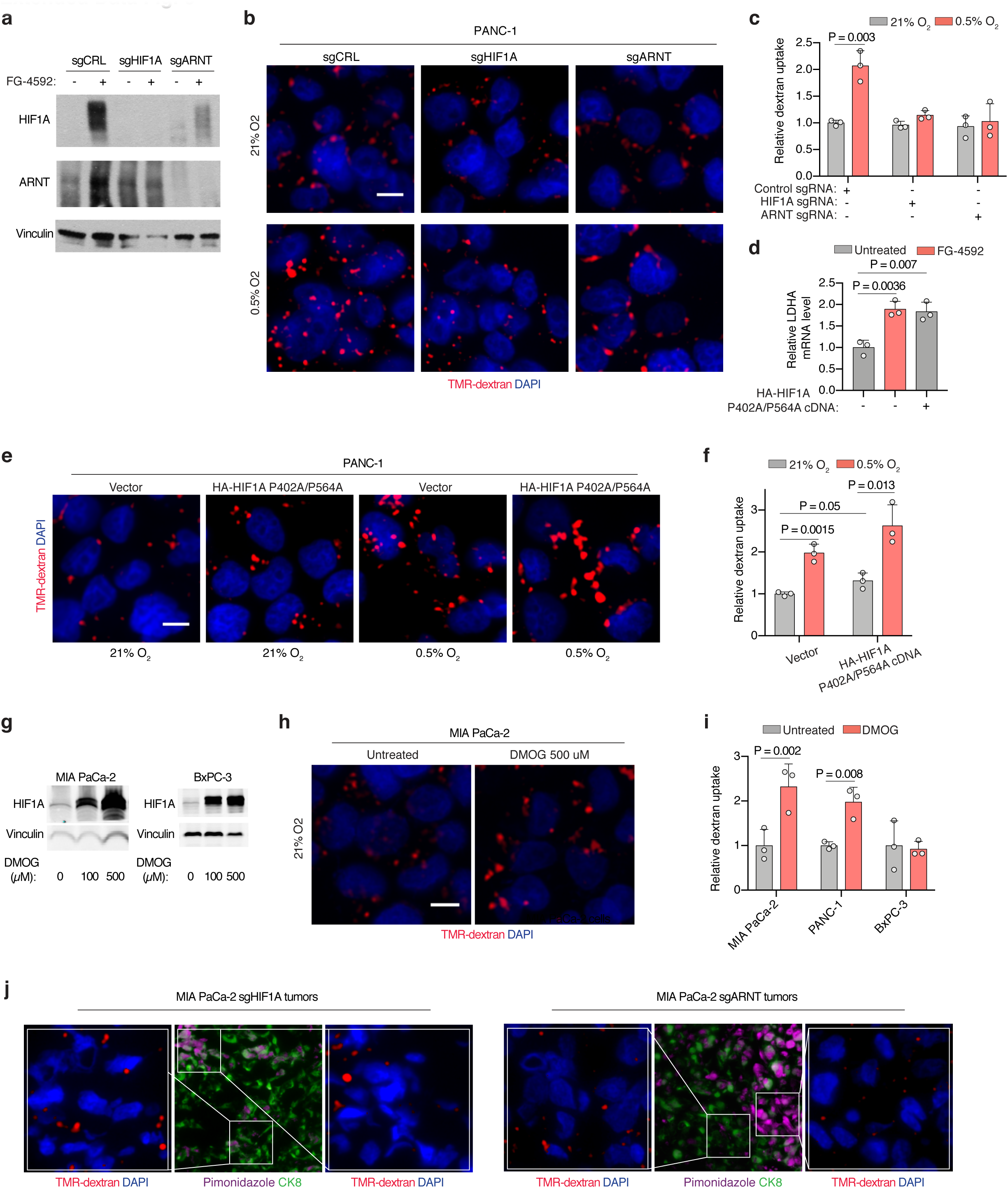
HIF1A stabilization stimulates macropinocytosis in PDAC cells and tumors. **(a)** Immunoblot analysis of HIF1A and ARNT in the indicated sgRNA-transduced PANC-1 cells treated with FG-4592 as shown. Vinculin was used as a loading control. **(b, c)** Representative images (**b**) and quantification (**c**) of TMR-dextran (red) uptake in control or sgHIF1A- or sgARNT-transduced PANC-1 cells under normoxia (21% O_2_) or hypoxia (0.5% O_2_). Nuclei are labeled with DAPI (blue). Scale bar, 10 μm. In **c**, data are presented relative to values obtained for control normoxic cells. **(d)** Relative mRNA levels of the HIF1A-target lactate dehydrogenase A (LDHA) in the indicated MIA PaCa-2 cell lines expressing a control vector or HA-HIF1A P402/P564A cDNA and treated with the prolyl-hydroxylase (PHD)-inhibitor FG-4592 (100 μM, 72 hrs) as shown. **(e, f)** Representative images (**e**) and quantification (**f**) of TMR-dextran (red) uptake in PANC-1 cells expressing a control vector or HA-HIF1A P402/P564A cDNA under normoxia (21% O_2_) or hypoxia (0.5% O_2_). Nuclei are labeled with DAPI (blue). Scale bar, 10 μm. In **f**, data are presented relative to values obtained for control normoxic cells. **(g)** Immunoblot analysis of HIF1A in MIA PaCa-2 and BxPC-3 cells treated with 0, 100, and 500 μM of DMOG under normoxia (21% O_2_). Vinculin was used as a loading control. **(h)** Representative images of TMR-dextran (red) uptake in MIA PaCa-2 cells in the absence or presence of DMOG (500 μM) under normoxia (21% O_2_). Nuclei are labeled with DAPI (blue). Scale bar, 10 μm. **(i)** Quantification of macropinocytic uptake in MIA PaCa-2, PANC-1 and BxPC-3 cells treated in the absence or presence of DMOG (500 μM) under normoxia (21% O_2_). Data are presented relative to values obtained for untreated cells. **(j)** Representative images of TMR-dextran (red) uptake from sections of xenograft tumors arising from MIA PaCa-2 cells transduced with sgRNAs targeting HIF1A or ARNT with tumor cells immunostained with anti-CK8 (green), and pimonidazole detected with anti-pimonidazole (purple). Nuclei are labeled with DAPI (blue). **c, d, f, i,** Bars represent mean ± s.e.m. At least 500 (**c, f, i**) cells were quantified in each biological replicate (*n* = 3). Statistical significance was determined by two-tailed unpaired *t*-test.

**Extended Data Figure 7.**
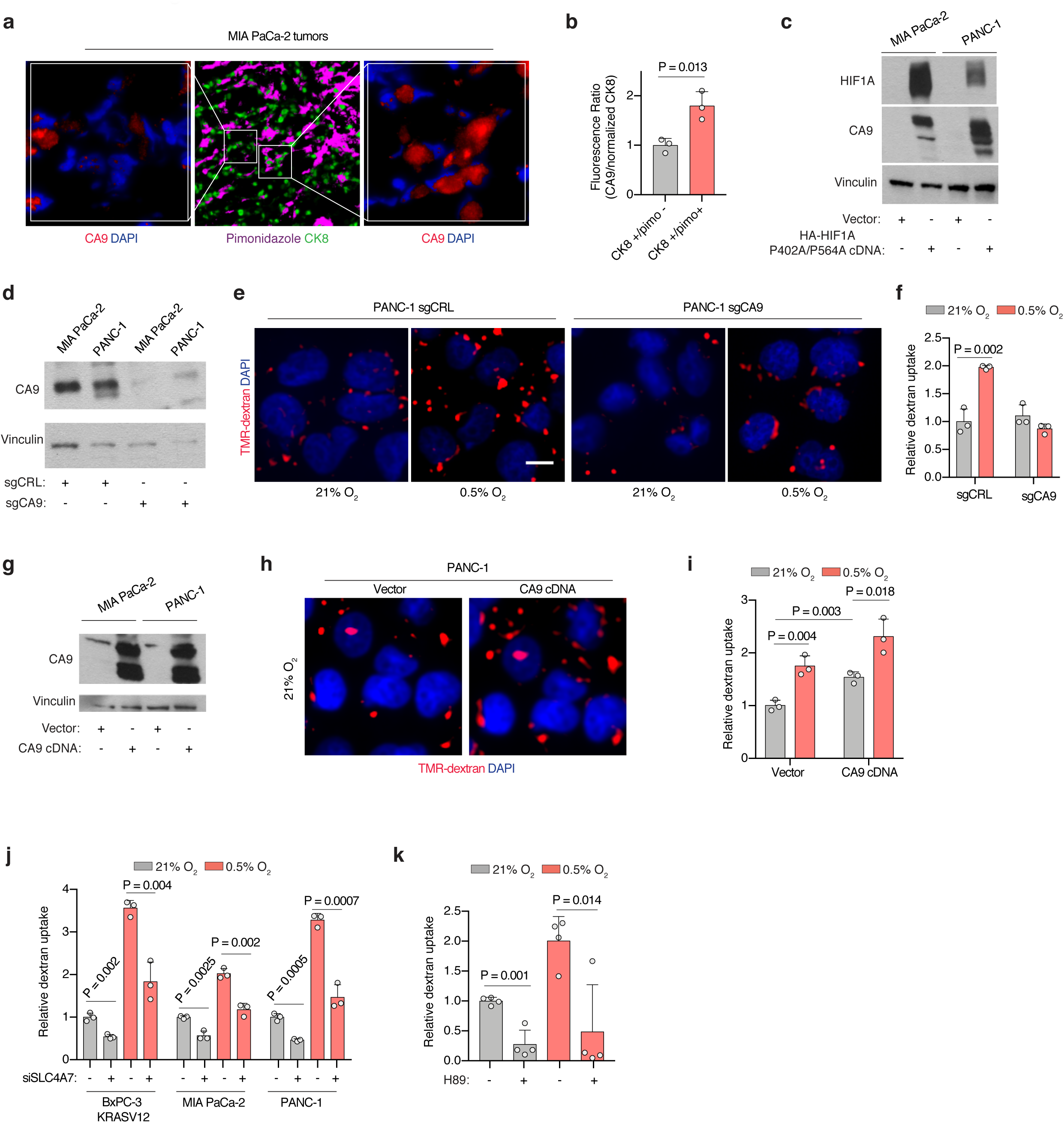
Bicarbonate generation by CA9 mediates the effect of HIF1A on PDAC macropinocytosis under hypoxia. **(a)** Representative images of CA9 expression (red) from sections of MIA PaCa-2 xenograft tumor tissue with tumor cells immunostained with anti-CK8 (green), and pimonidazole detected with anti-pimonidazole (purple). Nuclei are labeled with DAPI (blue). **(b)** Immunofluorescence of CA9 relative to CK8 in sections of xenograft tumors arising from MIA PaCa-2 parental cells. Data are presented relative to values obtained for CK8+/pimo-areas. **(c)** Immunoblot analysis of CA9 and HIF1A in indicated cell lines expressing a control vector or HA-tagged HIF1A P402A/P564A cDNA. Vinculin was used as a loading control. **(d)** Immunoblot analysis of CA9 in the indicated cell lines transduced with a control sgRNA or an sgRNA targeting CA9. Vinculin was used as a loading control. **(e, f)** Representative images (**e**) and quantification (**f**) of TMR-dextran (red) uptake in control or sgCA9-transduced PANC-1 cells under normoxia (21% O_2_) or hypoxia (0.5% O_2_). Nuclei are labeled with DAPI (blue). Scale bar, 10 μm. In **f**, data are presented relative to values obtained for control cells under normoxia. **(g)** Immunoblot analysis of CA9 in MIA PaCa-2 and PANC-1 cells transduced with CA9 cDNA or a control vector. Vinculin was used as a loading control. **(h)** Representative images of TMR-dextran (red) uptake in control or CA9 overexpressing PANC-1 cells under normoxia (21% O_2_) with macropinosomes labeled with TMR-dextran (red). Nuclei are labeled with DAPI (blue). **(i)** Quantification of TMR-dextran uptake in control or CA9 overexpressing PANC-1 cells under normoxia (21% O_2_) and hypoxia (0.5% O_2_). Data are presented relative to values obtained for control cells. **(j)** Quantification of TMR-dextran uptake in the indicated cell lines transfected with a control (-) or an SLC4A7 siRNA under normoxia (21% O_2_) and hypoxia (0.5% O_2_). Data are presented relative to values obtained for control normoxic cells. **(k)** Quantification of TMR-dextran uptake in MIA PaCa-2 cells treated with the PKA inhibitor H89 (15 μM) as shown under normoxia (21% O_2_) or hypoxia (0.5% O_2_). Data are presented relative to values obtained for control normoxic cells. **b, f, i, j, k,** Bars represent mean ± s.e.m. At least 300 (**b**) and 500 (**f, g, j, k, l**) cells were quantified in each biological replicate (*n* = 3). Statistical significance was determined by two-tailed unpaired *t*-test.

## MATERIALS AND METHODS

### Cell lines, compounds and constructs

All cell lines were obtained from American Type Culture Collection (ATCC), maintained under 5% CO_2_ at 37°C, and routinely tested for mycoplasma contamination every two months. HY15549 cell line is a gift from Nabeel Bardeesy lab and KP-Lung cell line is a gift from Thales Papagiannakopoulos. Human pancreatic stellate cells (hPSCs) were obtained from Dr. Rosa Hwang.

Antibodies to SLC1A3 (GTX20262) and Beta-Actin (GTX109639) were obtained from GeneTex; HRP-conjugated anti-rabbit antibody and ARNT antibody (sc-17811) from Santa Cruz; anti-FLAG M2 (F1804) and anti-T7 (69522) from Sigma; GOT1 antibody (ab170950) and total OXPHOS antibody cocktail (ab110411) from Abcam; GOT2 antibody from Invitrogen (PA5-27572); anti-Vinculin from CST (#4650); monoclonal antibody against HIF1A from BD Biosciences (#610958); anti-CA9 from Novus Biologicals (NB100-417); CK8 (MABT329) and MCT1 (AB3540P) antibodies from Millipore; and anti-pimonidazole from Hypoxyprobe (PAB2627AP).

DAPI was purchased from Vector Laboratories; sodium pyruvate, polybrene, puromycin, doxycycline hyclate and Ammonium Iron (III) Citrate (FAC) from Sigma; aspartic acid from Acros; blasticidin from Invivogen; antimycin A and piericidin from Enzo Life Sciences; uridine from Alfa Aesar; Matrigel from Corning; bafilomycin A1, DMOG, S0859, H89 and FG-4592 from Cayman Chemical; AR-C155858 was purchased from R&D Systems; pimonidazole hydrochloride from Hypoxyprobe (HP-1000); TMR-Dextran from Fina Biosolutions; and DQ Green BSA (D12050) from Thermo Fisher.

All cell lines were cultured in RPMI or DMEM medium containing 1 mM glutamine, 10% fetal bovine serum, penicillin and streptomycin. For isotope tracing experiments, [U-^13^C]-L-Glutamine (Cambridge, CLM-1822-H-0.1), RPMI 1640 without glutamine (Gibco, #21870076) and with dyalized serum (Gibco, #26400044) were used.

For generation of the lentiviral knockout constructs, annealed oligos (below) were cloned using a T4 ligase (NEB) into lentiCRISPR-v2 vector or into a lentiCRISPR-v1-GFP and lentiCRISPR-v1-RFP for generation of GOT1/GOT2 double knockout cell lines:

Mouse sgGOT1_F2, caccGAGGGGTGCAGTCTTTGGGA;
mouse sgGOT1_R2, aaacTCCCAAAGACTGCACCCCTC;
human sgGOT1_F4, caccGATGTGGTTCCAGGTCCCAG;
human sgGOT1_R4, aaacCTGGGACCTGGAACCACATC;
mouse sgGOT2_F2, caccGATGGGCGTGTGATTTCCCC;
mouse sgGOT2_R2, aaacGGGGAAATCACACGCCCATC;
human sgGOT2_F7, caccgATCTTTGCAGACCATAGTGA;
human sgGOT2_R7, aaacTCACTATGGTCTGCAAAGATc;
mouse sgMCT1_F, caccgAGCCTGATTAAGTGGAGCCA;
mouse sgMCT1_R, aaacTGGCTCCACTTAATCAGGCTc;
human sgHIF1A_1F, caccgAGGATGCTTGCCAAAAGAGG;
human sgHIF1A_1R, aaacCCTCTTTTGGCAAGCATCCTc;
human sgARNT_3F, caccgACTGGCAACACATCCACTGA;
human sgARNT_3R, aaacTCAGTGGATGTGTTGCCAGTc;
human sgCA9_9F, caccGTACCTGGCTCAGCAGCCCG;
human sgCA9_9R, aaacCGGGCTGCTGAGCCAGGTAC.

Same protocol was used to clone 5 sgRNAs used for competition sgRNA assays targeting control regions (sgINTERGENIC) or aspartate-malate shuttle genes. The sgRNAs sequences for mouse and human genes are as follow:

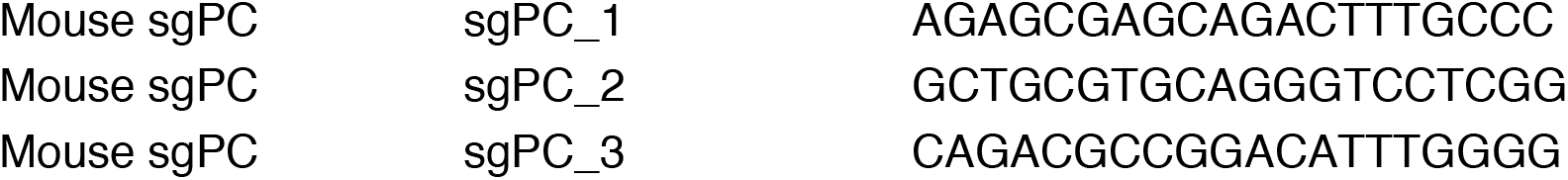
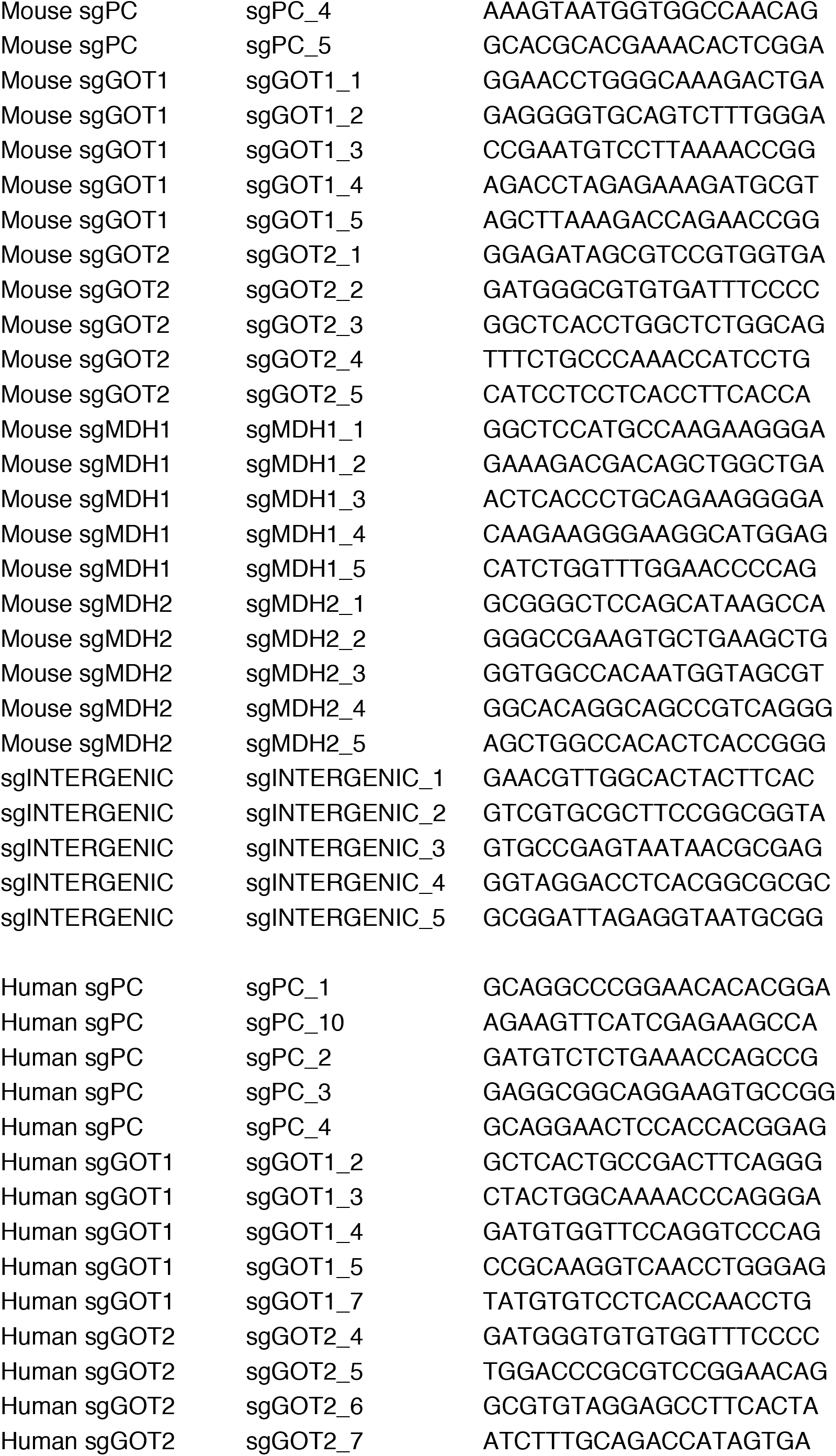
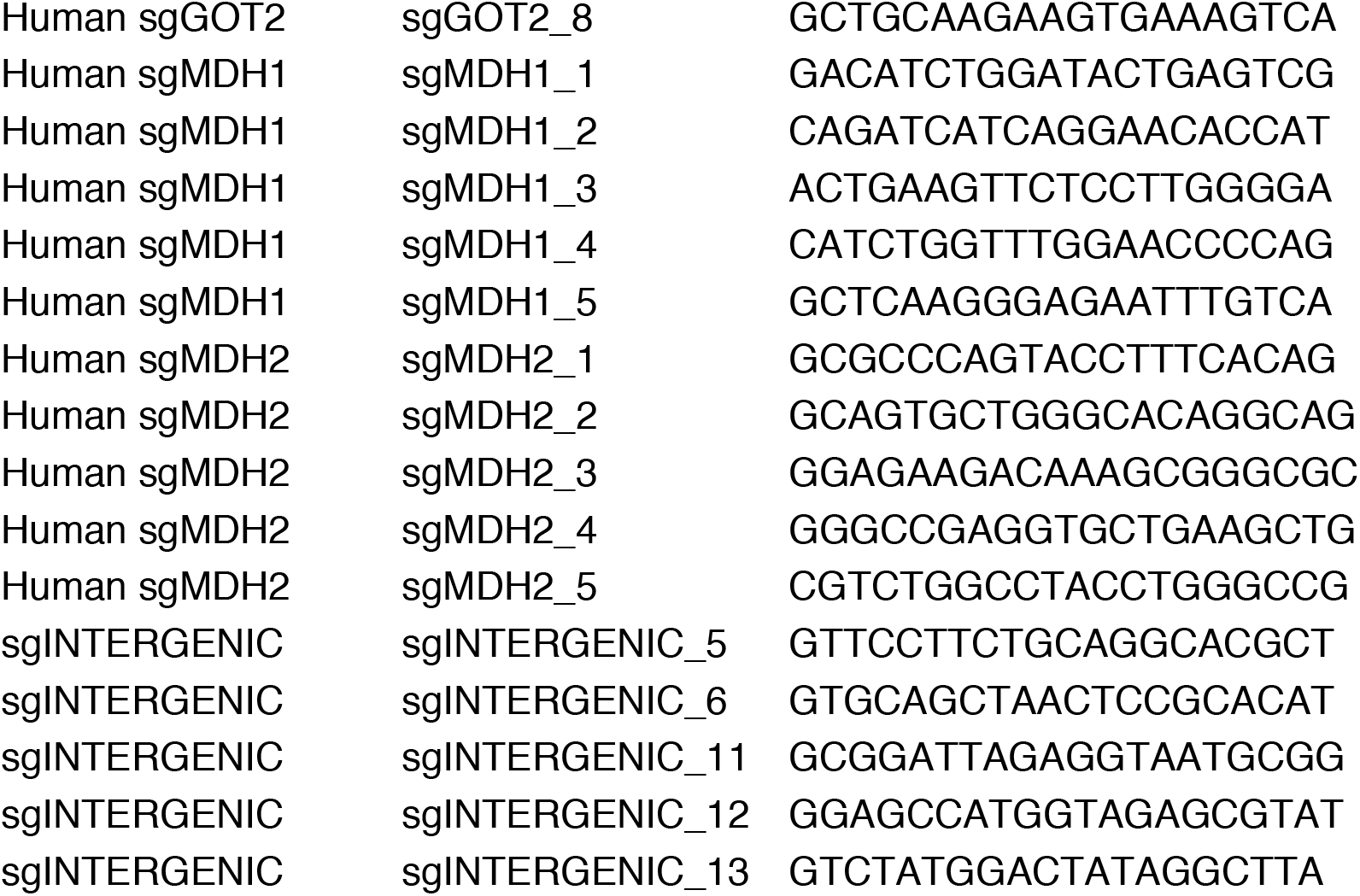

sgRNA-resistant GOT1 and GOT2 cDNAs; and cyto-lbNOX, mito-lbNOX, gpASNase and CA9 cDNAs were synthesized as gBlock fragments (IDT) and cloned into PMXS-IRES-Blast vector by Gibson assembly. MCT1 and SLC1A3 cDNAs cloned into PMXS-IRES-Blast were previously generated. Constitutively stable HIF1A construct was obtained from Addgene (HA-HIF1alpha P402A/P564A-pcDNA3)^37^, and amplified by PCR using the following primers prior to its cloning into PMXS-IRES-Blast: HIF1A_cDNA_F, GCCGGATCTAGCTAGTTAATTAAGCCACCATGGCCTACCCCTACGACGTGCCCGACT; HIF1A_cDNA_R, GGGCGGAATTTACGTAGCTCAGTTAACTTGATCCAAAGCTCTGAGTAATTC.

### Generation of Knockout and cDNA Overexpression Cell Lines

For generation of single gene knockout cell lines, lentiviral vector expressing the corresponding sgRNA was transfected into HEK293T cells with lentiviral packaging vectors VSV-G and Delta-VPR using XtremeGene transfection reagent (Roche). After 24 hr, media was aspirated and replaced by fresh media. 48 hr after transfection, virus particles contained in the supernatant were collected and filtered using a 0.45 mm filter to avoid collection of cells. Cells to be transduced were plated in 6-well tissue culture plates and infected in media containing the virus and 8 mg/ml of polybrene. To increase transduction efficiency, a spin infection was performed by centrifugation at 2,200 rpm for 1.5 hr. 48 hr after the infection, selection of transduced cells was initiated by addition of puromycin. For generation of single knockout cells for GOT1 and GOT2 HY15549 and MIA PaCa-2 lines, cells were raised from a single clone isolated by serial dilution of cells into a 96-well plate containing 0.1 mL of media. Single cell clones were grown for three weeks, and the resultant clones were validated by western blot and expanded. GOT2 knockout cells were generated, expanded and always maintained in RPMI media containing 1 mM pyruvate, except during proliferation experiments, in which low pyruvate media (50 uM) was used. This is due to a requirement of GOT2 KO cells for pyruvate.

For generation of GOT1/2 double knock-outs, cells were transduced with lentiviral GFP-sgGOT2 and next day with lentiviral RFP-sgGOT1. Double GFP/RFP positive cells were then single cell sorted using a BD FACSAriaII and expanded for 3 weeks in high aspartate RPMI media (20 mM Aspartate). Absence of GOT2 and GOT1 was validated by western blot and glutamine isotope labeling experiments. MIA PaCa-2 and PANC-1 HIF1A, ARNT and CA9 knockout cell lines were generated by the same procedure of puromycin selection but, instead of using expanded single cell clones, mixed cell populations were used.

For overexpression of cDNAs, retroviral vectors with indicated cDNAs were transfected with retroviral packaging plasmids Gag-pol and VSV-G into HEK293T cells. For virus collection and transduction, we followed the same procedure than that used with lentiviral vectors. Selection of transduced cells was achieved by addition of blasticidin.

For generation of MIA PaCa-2 Rho(0) cells, parental cells were maintained and passaged at low confluency in RPMI media containing pyruvate (1 mM) and uridine (100 uM) in the presence of ethidium bromide (1:10,000) for 30 days.

HeLa and BxPC-3 stable cell lines with doxycycline-inducible expression of FLAG-KRAS(G12V) or T7-KRAS(G12V) were generated with lentiviral particles. Cells were transduced with lentiviral particles containing pTRIPZ-FLAG-K-Ras(G12V), selected with puromycin (2 ug/mL) for 72 h, and maintained in 10% tetracycline-free FBS (Clontech).

### Proliferation Assays

Cell lines were plated in 6-well plates at 1,000-2,500 (HY15549) or 15,000 (MIA PaCa-2 and PA-TU-8898-T) cells per well in standard conditions. Next day, cells were treated with ETC inhibitors (Antimycin A or piericidin), left untreated in a standard incubator (normoxia) or cultured in a hypoxia chamber (INVIVO) set to 0.5% O_2_. For hypoxic conditions, culture media were pre-incubated for 24 hrs and added inside the hypoxia chamber to the corresponding cells. After 5-6 days of growth, cells were trypsinized and counted using a Beckman Z2 Coulter Counter with a size selection setting of 8–30 mm. All experiments with HY15549 GOT2_KO cells were performed in the presence of physiological levels of pyruvate (50 uM).

Experiments using conditioned media from human pancreatic stellate cells (hPSCs) were performed as previously described^23^. Briefly, hPSCs were maintained in RPMI media and plated in 10-cm dishes at 1×10^6^ cells/plate. Next day, media was changed and 10 mLs of RPMI were added. After 48 hrs, conditioned media was collected and filtered using a 0.45 mm filter to avoid collection of hPSCs.

For most of BSA rescue experiments Albumin, Bovine Serum, Fraction V, Fatty Acid-Free, Nuclease- and Protease-Free (Sigma, #126602) was used. Cells were plated in 6-well plates in media without BSA. Next day, media was aspirated and fresh media with or without BSA added (3 mLs/well), containing or not ETC inhibitors, or upon its pre-incubation overnight in the hypoxia chamber at 0.5% O_2_.

For BSA rescue experiments of CA9 knockout cell lines under hypoxia, cells were seeded at a density of 1,000-1,500 cells per well of a 96-well plate and incubated under normoxia for 24 hours in complete media. Media was then changed to complete media supplemented with 0.5% Fraction V BSA (Calbiochem). Cells were incubated for 5 days under either normoxia or hypoxia (0.5% O_2_). Cells were then fixed in 3.7% formaldehyde, washed, and incubated in a SYTO™ 60 red fluorescent nucleic acid stain (1:5000 dilution in 0.1% Triton/PBS) for 30 minutes at room temperature. Plates were scanned and imaged using a Licor Odyssey Classic imager and the Odyssey 2.1 software.

For induction of oncogenic KRAS expression prior to proliferation experiments in BxPC-3 with TET-inducible FLAG-KRAS(G12V), cells were treated with doxycycline (1 ug/mL) for 48 h prior to plating 20,000 cells/well in the presence of 0.1 ug/mL of doxycycline for 7 days. Doxycycline was added fresh every 3 days.

Micrograph pictures of endpoint in proliferation experiments were taken with Primovert microscope (Zeiss).

### Metabolite Profiling and Isotope Tracing

For the initial metabolite profiling experiment in KP PDAC cell lines lines, each indicated cell line (50,000 cells per replicate for HY15549; 100,000 cells per replicate for MIA PaCa-2 and AsPC-1) was cultured as triplicates in 6-well plates. After 12 hrs, cells were left untreated at normoxia (21% O_2_), treated for 20 hours with antimycin A (HY15549: 30 nM; AsPC-1: 50 nM; MIA PaCa-2: 50 nM) or grown in a hypoxia chamber at 0.5% O_2_ with media pre-incubated in the chamber for 24 hrs.

For glutamine tracing experiments in HY15549 and MIA PaCa-2 parental and GOT1/2 DKO lines, 50,000-100,000 cells per well were plated in RPMI media containing 20 mM aspartate. After 24 hrs, wells were thoroughly washed with PBS five times to remove all the aspartate, prior to addition of RPMI with dialyzed FBS and no aspartate. After additional 24 hrs, media was changed and replaced with 2 mL of custom RPMI media lacking glutamine and supplemented with [U-^13^C]-Glutamine (500 uM). Cells were cultured in this media for additional 8 hrs before their collection and isotope labeling analysis of aspartate and citrate.

For metabolite profiling under hypoxia or ETC inhibition in the presence or absence of BSA, MIA PaCa-2 parental cells were plated at low confluency (200,000 cells in 10-cm dishes in triplicates) in regular RPMI media. For hypoxia conditions, media with or without BSA (2% BSA) was pre-incubated in the hypoxia chamber overnight. Next day, media was aspirated and fresh normoxic or hypoxia media with and without BSA were added. After 24 hrs under these culture conditions, metabolite extraction was performed. For ETC inhibition profiling, same procedure was used but adding piericidin (150 nM) to each condition for 24 hrs.

For metabolite profiling of PDAC cells auxotroph for aspartate upon BSA addition, MIA PaCa-2 GOT1/2 DKO cells were plated at low confluency (200,000 cells in 10-cm dishes) in either RPMI media or RPMI media with BSA 2%, both prepared with dialyzed FBS to control for the BSA present in serum. After 48 hrs under these culture conditions, samples were collected, and metabolite analysis was performed.

In all experiments, at the time of collection cells were washed three times with 1-3 mL of cold 0.9% NaCl, and polar metabolites extracted in 1 mL of cold 80% methanol containing internal standards (MSK-A2-1.2, Cambridge Isotope Laboratories, Inc.) by scraping the cells in each well on ice and collecting the content in 1.5 mL tubes. After 10 min extraction by vortexing and centrifugation for 10 min at 10,000 × g and 4°C, samples were nitrogen-dried and stored at −80C until analysis by LCMS. Analysis was conducted on a QExactive benchtop orbitrap mass spectrometer equipped with an Ion Max source and a HESI II probe, which was coupled to a Dionex UltiMate 3000 UPLC system (Thermo Fisher Scientific, San Jose, CA). External mass calibration was performed using the standard calibration mixture every 7 days.

Dried polar samples were resuspended in 100 μL water and 2 μL were injected into a ZIC-pHILIC 150 x 2.1 mm (5 μm particle size) column (EMD Millipore). Chromatographic separation was achieved using the following conditions: Buffer A was 20 mM ammonium carbonate, 0.1% ammonium hydroxide; buffer B was acetonitrile. The column oven and autosampler tray were held at 25°C and 4°C, respectively. The chromatographic gradient was run at a flow rate of 0.150 mL/min as follows: 0–20 min.: linear gradient from 80% to 20% B; 20–20.5 min.: linear gradient from 20% to 80% B; 20.5–28 min.: hold at 80% B. The mass spectrometer was operated in full-scan, polarity switching mode with the spray voltage set to 3.0 kV, the heated capillary held at 275°C, and the HESI probe held at 350°C. The sheath gas flow was set to 40 units, the auxiliary gas flow was set to 15 units, and the sweep gas flow was set to 1 unit. The MS data acquisition was performed in a range of 70–1000 m/z, with the resolution set at 70,000, the AGC target at 10e6, and the maximum injection time at 20 msec. Relative quantitation of polar metabolites was performed with XCalibur QuanBrowser 2.2 (Thermo Fisher Scientific) using a 5 ppm mass tolerance and referencing an in-house library of chemical standards. For stable isotope tracing studies, fractional labeling was corrected for natural abundance using an in-house algorithm^50^.

Metabolite levels were normalized to the total protein amount for each condition.

Principle component analysis was performed on whole-cell metabolite profiles of three PDAC cell lines under normoxic, antimycin A and hypoxic conditions. The PCA is centered and scaled relative to the samples.

### Immunoblotting

Cell pellets were washed twice with ice-cold PBS prior to lysis in lysis buffer (10 mM Tris-Cl pH 7.5, 150 NaCl, 1 mM EDTA, 1% Triton X-100, 2% SDS, 0.1% CHAPS) supplemented with protease inhibitors (Roche). After sonication of each cell lysate and centrifugation for 10 min at 4C and 20,000 g, supernatants were collected, and their protein concentration were determined by using Pierce BCA Protein Assay Kit (Thermo Scientific) with bovine serum albumin as a protein standard. Depending on the experiment, samples were resolved on 8%, 12%, 12-16% or 16% SDS-PAGE gels and transferred to Immobilon-P PVDF membranes (Millipore) with 0.45 um (CA9) or 0.2 um (all the other proteins) pore size. Membranes were blocked in 5% milk (for analysis of GOT1, GOT2, ACTB, FLAG, T7, SLC1A3, MCT1 and total OXPHOS) or BSA 5% (for HIF1A, ARNT, CA9 and vinculin) and analyzed by immunoblotting as previously described.

### Mouse studies

All animal studies and procedures were conducted according to a protocol approved by the Institutional Animal Care and Use Committee (IACUC) at the Rockefeller University and by the New York University Grossman School of Medicine Institutional Animal Care (IACUC). All mice were maintained on a standard light-dark cycle with food and water ad libitum.

Xenograft tumors were initiated by injecting the following number of cells in DMEM with 30% Matrigel:

0.5 million cells/100 uL for mouse HY15549 parental and individual GOT knock out cell lines.
3 million cells/100 uL for MIA PaCa-2 parental and individual GOT knock out cell lines.
5 milllion cells/100 uL for HY15549 and MIA PaCa-2 GOT1/2 DKO cell lines.

After injections in the left and right flanks of male and female 6-9 weeks old NOD scid gamma (NSG) mice (Taconic), tumors were grown for 3-6 weeks.

For orthotopic pancreas injections, we followed previously described protocols^51^. Briefly, a small incision was made on the upper left quadrant of the abdomen and the pancreas was externalized. 100,000 cells in 50 uL of PBS with 50% Matrigel were injected into the pancreatic tail with insulin syringes (29-gauge needle, BD).

For metabolite profiling of MIA PaCa-2 DKO tumors, animals were sacrificed and tumors were extracted and immediately lysed in cold 80% methanol containing internal standards by using a Bead Ruptor 24 (Omni International) by 6 cycles of 20 s at 6 m/s. Supernatants were collected after 10 min centrifugation at 10,000 × g and 4 C, nitrogen dried and analyzed for the content of carbon labeled aspartate.

### Real Time PCR assays

RNA was isolated by an RNeasy Kit (Qiagen) according to the manufacturer’s protocol and samples were analyzed as previously described^52^. Briefly, RNA was spectrophotometrically quantified and equal amounts were used for cDNA synthesis with the Superscript II RT Kit (Invitrogen). qPCR analysis was performed on an ABI Real Time PCR System (Applied Biosystems) with the SYBR green Mastermix (Applied Biosystems).

The primers used were the following: LDHA_F, ATGGCAACTCTAAAGGATCAGC; LDHA_R, CCAACCCCAACAACTGTAATCT; ACTINB_F, TTTTGGCTATACCCTACTGGCA; ACTINB_R, CTGCACAGTCGTCAGCATATC.

Results were normalized to ACTINB.

### CRISPR-based screen

The highly focused metabolism sgRNA library used for CRISPR screens in HY15549 was generated as previously described and contains sgRNAs targeting the mouse genes corresponding to our previously described highly focused metabolism human library^53^. Briefly, the plasmid pool was used to generate a lentiviral library containing 5 sgRNAs per gene target. This viral supernatant was titered in HY15549 cells by infecting target cells at increasing amounts of virus in the presence of polybrene (8 μg/ml) and by determination of cell survival after 3 days of selection with puromycin. 2 × 10^6^ of each cell type were infected at a MOI of 1 prior to selection with puromycin for 3 days. An initial pool of 2 million cells was harvested. Infected cells were then cultured for 14 population doublings at 21% O_2_, 0.5% O_2_ or in the presence of an ETC inhibitor, piericidin (40 nM). After this, 2 million cells were collected, and their genomic DNA extracted by a DNeasy Blood & Tissue kit (QIAGEN). For amplification of sgRNA inserts, we performed PCR using specific primers for each condition. PCR amplicons were then purified and sequenced on a NextSeq 500 (Illumina). Sequencing reads were mapped and the abundance of each sgRNA was measured. Gene score is defined as the median log_2_ fold change in the abundance between the initial and final population of all sgRNAs targeting that gene. A gene score lower than −1 is considered as significantly essential.

### sgRNA competition assay *in vitro* and *in vivo*

Five control sgRNAs targeting intergenic regions, and five sgRNAs targeting *GOT1*, *GOT2*, *MDH1*, *MDH2* or *PC* were cloned into linearized lentiCRISPR-v2 vector and transformed in NEB competent E. coli. Each plasmid was then pooled at equal concentrations and used for lentivirus production as previously described. Each cell line was infected and selected with puromycin for 3 days prior to being *in vitro* cultured or, in the case of PANC-1, also injected subcutaneously or orthotopically in the pancreas of NSG mice. Tumors were collected after 2-4 weeks of growth. An initial pool of each sample was taken for normalization. After 14-21 days gDNAs were isolated and amplified by PCR. PCR amplicons were then sequenced on a NextSeq 500 (Illumina). Guide scores were calculated as median log_2_ fold change in the abundance between the initial and final population of that sgRNA similar to standard CRISPR screens. The KP PDAC patient derived xenograft (PDX) model for *in vivo* competition assay was described previously^51^. Briefly, low passage PDX pancreatic tumor was chopped finely and digested with collagenase type IV for 30 min. Next, mouse cells were removed from the single-cell suspension via magnetic-associated cell sorting using the Mouse Cell Depletion Kit (130-104-694, Miltenyi), resulting in a single-cell suspension of predominantly pancreatic cancer cells of human origin. 10 million cells of this PDX suspension were transduced with the human aspartate-malate shuttle lentiviral particles and subsequently injected subcutaneously in PBS with 50% Matrigel in immunodeficient mice 24 hrs after infection. PDX tumors were collected after 3 weeks of growth, their genomic DNA isolated and amplicons were amplified by PCR and sequenced.

### *In vitro* Macropinocytosis Assay

Cells were seeded onto glass coverslips in a 24 well plate. Forty-eight hours after seeding, cells were subjected to serum starvation. For 24 hour hypoxia experiments, cells were grown in normoxia (21% O_2_) for 24 hours, then incubated in a hypoxia chamber set at <1% O_2_-Whitley H35 Hypoxystation (Don Whitley Scientific) - for 24 hours prior to the assay. Macropinosomes were quantified as previously described^8^. Cells were incubated for 30 minutes in 70-kDa high-molecular weight TMR-dextran (Fina Biosolutions) diluted to a concentration of 1 mg/mL in serum-free media. Cells were then washed with ice-cold PBS and fixed with 3.7% formaldehyde. Cells were DAPI stained for nuclei and mounted onto glass slides with DAKO mounting medium (Agilent). Images were captured at 20x magnification using a Nikon Eclipse Ti2 microscope and analyzed using the ImageJ software (NIH). Background subtraction was applied, a threshold was set to detect macropinosomes, and a macropinocytic index was determined by the total particle area per cell.

### *In vivo* Macropinocytosis Assay

2 million cells of corresponding MIA PaCa-2 cell lines were resuspended in 50% Matrigel (Corning) and injected subcutaneously into the flanks of 7 week old female nude mice (Taconic). When tumors attained a volume of 500 mm^3^, pimonidazole (Hypoxyprobe, pimonidazole hydrochloride, HP-1000) was used to assess hypoxia *in vivo,* and TMR-dextran was used to label macropinosomes *in vivo* as previously described^8^. 1.5 mg pimonidazole dissolved in PBS was injected intraperitoneally using an insulin syringe. 100 uL of 10 mg/mL TMR-dextran was then injected intratumorally using an insulin syringe. Both pimonidazole and TMR-dextran were allowed to circulate for 90 minutes prior to euthanization. Mice were euthanized with CO_2_, and cervical dislocation as a secondary method according to approved IACUC protocols. Tumors were harvested and rapidly frozen in Tissue-Tek O.C.T. Compound. Frozen tumors were sectioned and stained for CK8, pimonidazole, and DAPI. Macropinocytosis was quantified as previously described^8^.

### *In vitro* DQ-BSA Assay

Cells were seeded onto glass coverslips in a 24 well plate. Forty-eight hours after seeding, cells subjected to serum starvation. Cells were incubated for 30 minutes in DQ Green BSA diluted to 0.05 mg/mL in serum-free media, then chased with serum-free media free of DQ BSA for 1.5 hours. Cells were then washed with ice-cold PBS and fixed with 3.7% formaldehyde. Cells were DAPI stained for nuclei and mounted onto glass slides with DAKO mounting medium (Agilent). Images were captured at 20x magnification using a Nikon Eclipse Ti2 microscope and analyzed using the ImageJ software (NIH). Background subtraction was applied, and a threshold was set to detect DQ BSA fluorescence.

### Immunofluorescence

Frozen tumor sections on glass slides or cells seeded on glass coverslips were washed in PBS and fixed in 3.7% formaldehyde for 15 minutes at room temperature. Non-specific signal blocking was done using 10% goat serum and 2% BSA in PBS for 1 hour at room temperature for tumor sections. Blocking in *in vitro* samples included a permeabilization step and was done using Triton X-100 (1:1000), 1% goat serum, and 3% BSA in PBS for 1 hour at room temperature. After incubation in primary antibody for 2 hours at room temperature, samples were washed and incubated with fluorescence-conjugated secondary antibodies. Images were captured at 20x or 60x magnification using the Nikon Eclipse microscope and analyzed using the ImageJ software (NIH). Hypoxic areas of tumor sections were identified by setting a threshold for pimonidazole fluorescence. The macropinocytic index within areas of high pimonidazole and lower pimonidazole was calculated by dividing the total area of macropinosomes by the total area of CK8 positive tumor cells. The following primary antibodies were used: CK8 (1:180) [Millipore MABT329], pimonidazole (1:20), and CA9 (1:100). The following secondary antibodies were used: Alexa Fluor™ 647 goat anti-rabbit and Alexa Fluor™ 488 goat anti-rat (1:500).

#### Statistics and reproducibility

GraphPad PRISM 7 and Microsoft Excel 15.21.1 software were used for statistical analysis. XCalibur QuanBrowser 2.2 (Thermo Fisher Scientific) was used for metabolomic analyses and ImageJ (NIH) for image analysis. Error bars, *P* values and statistical tests are reported in the figure captions. All experiments (except metabolite profiling experiments and *in vivo* tumor dextran assays, which were done once) were performed at least two times with similar results. Both technical and biological replicates were reliably reproduced.

#### Data availability

All data supporting the findings of this study are available from the corresponding authors on reasonable request.

